# Dispersal transiently modifies the temperature dependence of ecosystem productivity after an extreme thermal fluctuation

**DOI:** 10.64898/2025.12.29.695627

**Authors:** Keila Stark, Patrick L. Thompson, Joey R. Bernhardt, Coreen Forbes, Kaleigh E. Davis, Evgeniya Yangel, Andrea S. Jackman, Malena Heinrichs, Laura Wegener Parfrey, Mary I. O’Connor

## Abstract

Effects of warming on ecosystem productivity are typically summarized over broad time scales, yet they emerge from communities that can reorganize in a matter of days. Temperature accelerates ecosystem productivity through predictable effects on metabolic rates, but dispersal across thermally heterogeneous metacommunities may modify this effect by redistributing communities and their underlying thermal phenotypes. Using multi-trophic freshwater mesocosm communities in which warming and dispersal were manipulated, we tested the hypothesis that increasing dispersal modifies the thermal sensitivity of gross primary productivity (GPP) through the redistribution of phytoplankton biomass, size spectra, and thermal phenotype composition along spatial thermal gradients. High dispersal temporarily weakened the thermal sensitivity of GPP after an unplanned heatwave. This effect was caused by reduced mass-specific GPP in the warmest mesocosms accompanied by regional homogenization of phytoplankton communities and the spread of poorly adapted thermal phenotypes under the highest dispersal treatments. Except immediately post-heatwave, the thermal sensitivity of GPP was robust to dispersal, despite evidence of metacommunity dynamics. These findings suggest that the temperature dependence of ecosystem metabolism can be briefly modified by dispersal after a perturbation, but remains robust under steady-state conditions.

## Introduction

Temperature regulates ecosystem productivity through predictable kinetic effects on individual photosynthesis and respiration rates (Allen *et al*., 2005; Bardgett *et al*., 2008; Falkowski *et al*., 2000). Metabolic scaling theory predicts the thermal sensitivity (also known as temperature dependence or activation energy, *E*) of ecosystem productivity by scaling the thermal sensitivity of individual metabolism to whole communities when the biomass, body size distribution, and average thermal phenotype of constituent organisms are known (Allen *et al*., 2005; Enquist *et al*., 2003; Kerkhoff *et al*., 2005; Schramski *et al*., 2015). By assuming that these properties underlying whole-community metabolism are at steady-state, metabolic scaling approximates the thermal sensitivity of ecosystem productivity averaged across broad spatiotemporal scales (e.g., annual; Yvon-Durocher *et al*. 2012). At finer scales, however, the thermal sensitivity of metabolism from individuals to ecosystems can change rapidly with resource dynamics (Vinton & Vasseur, 2022), plasticity and adaptation (Kordas *et al*., 2022b; Padfield *et al*., 2016; Schaum *et al*., 2017), species interactions (Cruz *et al*., 2023; Davis *et al*., 2025; Garcia *et al*., 2023), and exchange of resources with other ecosystems (Yvon-Durocher *et al*., 2012). Therefore, the instantaneous kinetic effects of warming on metabolism may in turn affect ecological dynamics, restructuring the community composition and biomass underlying ecosystem productivity on intermediate time scales. Understanding how these dynamics modify metabolic scaling relationships may clarify ecosystem responses to increasingly unpredictable temperature regimes (O’Connor *et al*., 2025; Stark *et al*., 2025).

Dispersal, or the exchange of individuals among spatially structured groups of communities (i.e., metacommunities; Leibold *et al*. 2004), interacts with temperature to influence community structure (de Boer *et al*., 2014; Grainger & Gilbert, 2017; Limberger *et al*., 2014; Parain *et al*., 2019; Thompson *et al*., 2015). Dispersal can maintain diversity through spatial insurance, where negative effects of environmental change are mitigated by immigration (Loreau *et al*., 2003; Thompson *et al*., 2020). When habitat patches differ suffciently in abiotic conditions, moderate dispersal is thought to allow phenotypes to migrate into patches with optimal conditions for growth and reproduction (Chase & Leibold, 2003; Leibold *et al*., 2004; Tilman, 1982). Systems experiencing very low dispersal rates (i.e., dispersal-limited) tend to be more susceptible to extirpation from environmental perturbations due to a lower likelihood of demographic rescue (France & Duffy, 2006; Melbourne & Hastings, 2009; Mori *et al*., 2018). Excessively high dispersal is associated with mass effects, which homogenize compositional differences among communities and maintain phenotypes in suboptimal habitats where they might otherwise not persist (Leibold *et al*., 2004; Thompson *et al*., 2020). Spatial insurance facilitated by dispersal can enhance ecosystem productivity (Leibold *et al*., 2017; Thompson & Gonzalez, 2016; Tilman *et al*., 2014; Venail *et al*., 2010), however support for this effect varies across trophic levels and environments (Eggers *et al*., 2012; Limberger *et al*., 2019; Symons & Arnott, 2013; Thompson & Shurin, 2012). In the context of metabolic scaling, it follows that dispersal and its consequences for community structure could modify the thermal sensitivity of ecosystem productivity. To date, evidence for the temperature scaling of ecosystem productivity comes from field and outdoor experimental systems that are considered open to dispersal, but of unknown magnitude (Barneche *et al*., 2021; O’Connor *et al*., 2011; Padfield *et al*., 2017; Yvon-Durocher & Allen, 2012; Yvon-Durocher *et al*., 2011). Absent manipulative tests, it is unclear whether the inferred thermal sensitivity of productivity in these studies would be upheld at lower or higher dispersal magnitudes.

Given this, we tested the following hypotheses: (H1) Different dispersal magnitudes modify the thermal sensitivity of gross primary productivity (GPP); (H2) Dispersal modifies GPP along a spatial thermal gradient by non-exclusively affecting three community properties: total biomass, size spectra, and thermal phenotype composition. To test how dispersal modifies the thermal sensitivity of GPP (henceforth *E*_GPP_, the slope of the temperature-GPP relationship), we consider how these community properties are predicted to differ along thermal gradients within metacommunities (Allen *et al*., 2005; Padfield *et al*., 2017; Yvon-Durocher & Allen, 2012), and how dispersal is predicted to redistribute them (Leibold *et al*., 2004; Loreau *et al*., 2003; Thompson *et al*., 2020).

First, dispersal may modify GPP by redistributing biomass among communities. All else equal, more photosynthetic biomass yields higher productivity. Different dispersal magnitudes may therefore maintain or remove temperature-driven differences in biomass among communities, strengthening or weakening *E*_GPP_ along a spatial thermal gradient (Fig. 1a). Studies in forests and streams spanning regional to global extents have found that the positive relationship between temperature and ecosystem productivity is attributed to higher standing biomass at warmer temperatures, such that mass-specific productivity is temperature-invariant (Michaletz *et al*., 2014; Padfield *et al*., 2017). Assuming these findings reflect moderate (replete) dispersal conditions, *E*_GPP_ along a spatial thermal gradient should be comparatively reduced in low dispersal (-limited) systems, which often show higher demographic stochasticity and a weaker influence of environmental conditions on community assembly and biomass (Chase, 2010; Ron *et al*., 2018; Thompson *et al*., 2020). In high dispersal systems, *E*_GPP_ is further reduced due to regional homogenization of biomass differences (Fig. 1a).

**Figure 1.**
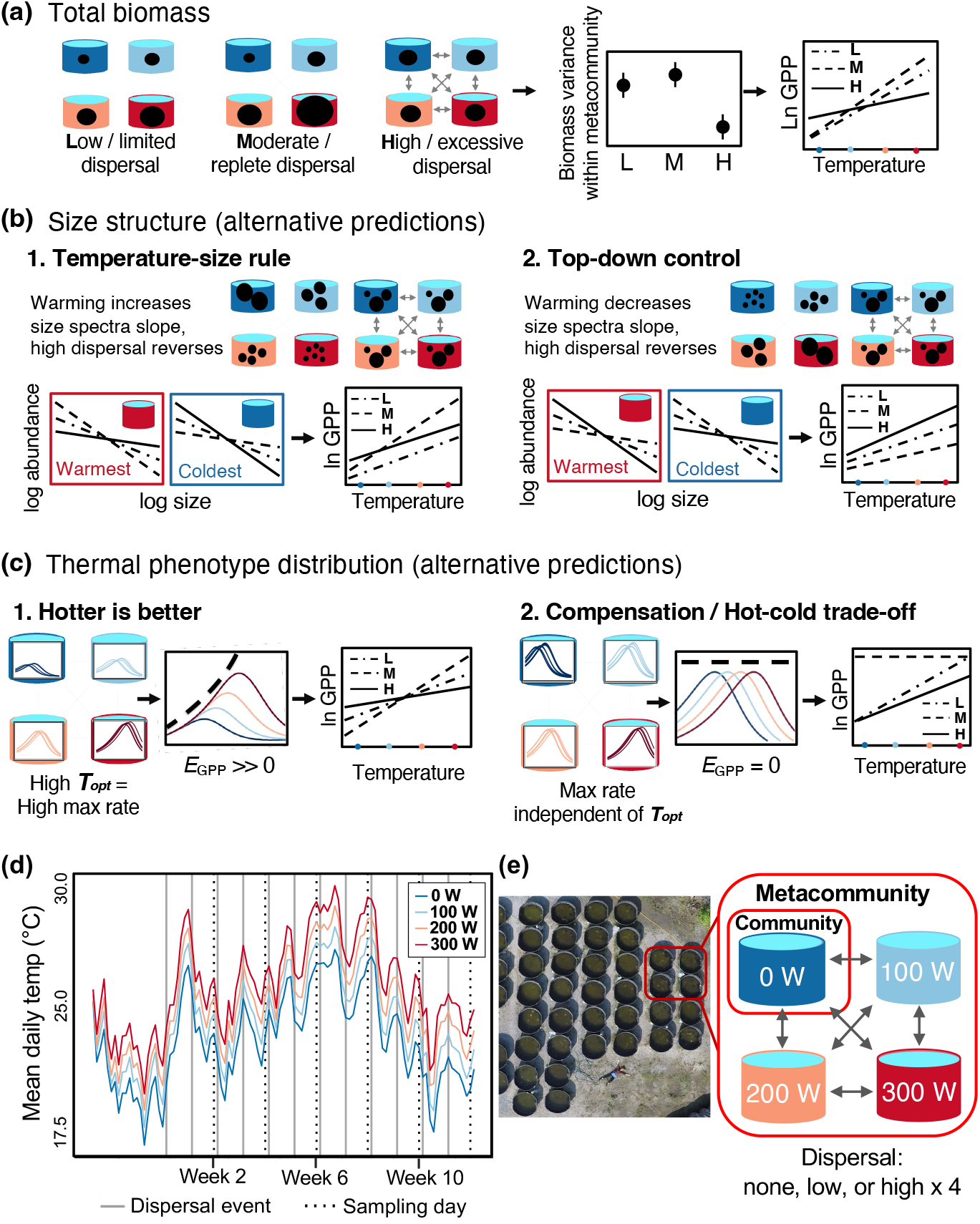
Predictions and experimental design. Mesocosm cartoons reflect the experimental design, with metacommunities comprised of four communities along a thermal gradient (tank colours) and linked by reciprocal dispersal. **(a)** Relative to low dispersal, temperature-driven differences in biomass are strengthened under moderate dispersal via species sorting and reduced under high dispersal via mass effects; this yields differences in biomass variance and therefore the thermal sensitivity of GPP (*E*_GPP_). **(b)** Higher dispersal may increase or decrease size spectra slopes depending on their response to warming (temperature-size rule versus top-down control) thereby increasing or decreasing *E*_GPP_ relative to a moderate dispersal scenario. **(c)** High dispersal increases or decreases *E*_GPP_ relative to moderate dispersal based on the relationship between thermal optima and maximum photosynthetic rate (hotter-is-better versus compensation). **(d)** The experiment time series begins on day 0 when communities were established. Coloured lines represent mean daily temperature in each warming (wattage) treatment. Grey solid lines show weekly dispersal treatment events. Black dotted lines show biweekly community sampling events. **(e)** Aerial view of the 48 mesocosms (researcher for scale). Inset diagram summarizes warming and dispersal treatments.

Second, dispersal may modify GPP by redistributing different-sized organisms, thereby modifying size spectra or relative abundances of small versus large organisms (Fig. 1b). Community size distribution affects ecosystem productivity because larger organisms generally have lower mass-specific metabolic rates according to sub-linear allometric scaling laws (Kleiber, 1947; West *et al*., 1997). Therefore, for the same total biomass and temperature, a community composed of large organisms is expected to have a lower total metabolic rate than one with smaller organisms (Barneche *et al*., 2014; Padfield *et al*., 2018; Yvon-Durocher & Allen, 2012). Temperature may affect size spectra in two ways. According to the temperature-size rule (TSR; Atkinson 1994), ectotherm body size is reduced at higher temperatures, which has been observed in phytoplankton laboratory monocultures (Bernhardt *et al*., 2018). In multi-trophic aquatic communities, however, warming can yield larger average phytoplankton sizes via intensified zooplankton grazing that targets smaller, easily-handled cells (Litchman *et al*., 2007; O’Connor, 2009; Padfield *et al*., 2018; Rall *et al*., 2009; Yvon-Durocher *et al*., 2015). We predict that temperature-driven differences in size spectra are most pronounced under moderate dispersal, and slightly reduced under low dispersal. In a TSR scenario, warmer communities have steeper size spectra (relatively more small individuals) and colder ones have shallower ones, strengthening *E*_GPP_ across a spatial thermal gradient because small organisms have higher mass-specific metabolism (Fig. 1b). Alternatively under top-down control, warming produces shallower size spectra, weakening *E*_GPP_ relative to a TSR scenario. In both cases, excessively high dispersal reverses these temperature-driven differences in size spectra, but in opposite directions (Fig. 1b).

Finally, dispersal may modify GPP by redistributing thermal traits or phenotypes, described by thermal response curves for metabolic or growth rate (Angilletta, 2009; Huey & Kingsolver, 1989; Thomas *et al*., 2012). A community’s thermal phenotype composition partially determines its productivity at a given temperature, such that *E*_GPP_ is sometimes taken to reflect spatial turnover in thermal phenotypes. For example, in Iceland’s Hengill geothermal field, mass-specific GPP was found to be temperature-independent across streams spanning a 40°C gradient (Padfield *et al*., 2017). Higher-than-expected photosynthetic rates in the coldest streams via cold-adaptation were responsible for this temperature-independence of GPP: an example of “metabolic compensation” or a “hot-cold trade-off” (Clarke, 1993; Padfield *et al*., 2017), where organisms modify their physiology to overcome temperature’s kinetic effects on photosynthetic enzymes (Atkin *et al*., 2015; Atkin & Tjoelker, 2003). Alternatively, there is substantial evidence of a “hotter-is-better” pattern wherein taxa with higher thermal optima also have higher maximum biological rates, such that thermal sensitivities across broad spatial or taxonomic extents are always positive (Angilletta *et al*., 2010; Eppley, 1972; Frazier *et al*., 2006; Kingsolver & Huey, 2008; Knies *et al*., 2009).

These patterns represent ends of a spectrum where complete compensation reflects plasticity or adaptation overcoming kinetic temperature effects, and hotter-is-better reflects adaptation within hard thermodynamic constraints (Kontopoulos *et al*., 2020). In spatially explicit comparisons, both patterns are contingent on phenotypic sorting into optimal thermal environments, which is facilitated by moderate dispersal according to metacommunity theory (Thompson *et al*., 2020). Under low and high dispersal, community thermal match may be dampened by stochastic loss or continual redistribution of well-adapted phenotypes, respectively. Both may result in a weaker *E*_GPP_ compared to moderate dispersal under a hotter-is-better scenario, or the opposite under compensation (Fig. 1c). We assess this hypothesis using a Community Thermal Phenotype Index (see Methods), which is similar to community weighted mean trait value in the trait-based ecology literature (De Bello *et al*., 2021).

Dispersal interacts with successional processes and thermal fluctuations to dynamically affect community structure and productivity. The geothermal stream example may reflect a dispersal-replete system because some streams are separated by a few metres and organisms disperse via wind, animals, and groundwater (Demars *et al*., 2011; Kordas *et al*., 2022a; O’Gorman *et al*., 2014). However, stream temperatures have been relatively constant over long timescales such that typical dispersal magnitudes that facilitate adaptation (Padfield *et al*., 2017) may not be sufficient in a hypothetical scenario where thermal fluctuations limit adaptation to one specific temperature. Thermal fluctuations can obscure or modify the spatial insurance effect (de Boer *et al*., 2014; Thompson *et al*., 2015) and cause a given dispersal magnitude to produce different effects on diversity across successional stages (Parain *et al*., 2019). We thus experimentally evaluated our hypotheses in the context of successional dynamics and thermal fluctuations by establishing freshwater communities in outdoor mesocosms exposed to ambient thermal fluctuations, including an unplanned heatwave mid-experiment.

## Materials and methods

### Experimental design and timeline

We assembled 48 mesocosms grouped into 12 metacommunities at the UBC experimental ponds facility in Vancouver, Canada (49.25, -123.23). Each community consisted of freshwater microbiota and invertebrates in 1136 L livestock watering tanks (Fig. 1e). Mesocosms were filled with water from an artificial pond nearby. We collected biota and sediment from five lakes within a 50 km radius using a 64 *μ*m zooplankton tow and Ekman grab, and transported them to the experiment site in large buckets. We homogenized the regional pool in one container and distributed it equally by volume among all mesocosms.

Each metacommunity consisted of four neighbouring communities (mesocosms) linked by the same dispersal treatment level: none (ambient), low, or high. Dispersal treatments entailed reciprocally exchanging equal volumes of water among all communities in a metacommunity (Fig. 1e). We rinsed equipment with tap water between treatments to minimize unintentional dispersal. In all dispersal treatments, we removed 30 L from each community, dividing each into three 10 L buckets. In the high-dispersal treatment, each community received 10 L from each of the other three communities in its metacommunity. In low-dispersal treatments, each community received 1 L from each other community, and the remaining 27 L that had been removed was returned. In no-dispersal treatments, we returned the full 30 L to its source community to standardize disturbance. These volume-based dispersal magnitudes roughly reflect dispersal rates in natural pond metacommunities (approximately 1% of total abundance, Michels *et al*. 2001).

We established spatial temperature gradients within each metacommunity using aquarium heaters (JÄGER TruTemp, EHEIM GmbH & Co KG, Germany) of different wattages (100 W, 200 W, 300 W) and dummy heaters (0 W). We used HOBO temperature loggers (Onset, USA) to record water temperatures hourly. The heaters imposed equal and consistent warming magnitudes of up to 3.0 °C (±0.4 °C) above ambient (Fig. 1d). Warming treatments were maintained as mesocosms were exposed to natural temperature fluctuations. Maximum temperatures were reached during the heatwave in week 7 (29.9 °C at 300 W) and minimum temperatures were reached shortly before the final sampling (18.7 °C at 0 W).

Mesocosms were seeded and exposed to warming treatments on May 25 2018 (day 0). We applied the first dispersal treatment on day 20. On day 18, we added sodium nitrate (109 *μ*g L^−1^) and potassium phosphate (15 *μ*g L^−1^) to each mesocosm to enhance productivity. Phosphate and nitrate levels did not differ significantly among experimental treatments (Fig. S1). The first sampling occurred on day 33 and roughly every two weeks thereafter (Fig. 1d) until the last sampling on September 5 (day 103). For simplicity, we refer to sampling days as the number of weeks after the first dispersal treatment (2, 4, 6, 8, 10, 12). We sampled GPP and phytoplankton communities (biomass, size spectra, taxonomic composition) on all six sampling days (Fig. 1d), zooplankton communities (biomass and taxonomic composition) in weeks 4, 8, and 12, and bacterial communities (taxonomic composition) in weeks 4 and 12 only.

### Estimating GPP

We estimated GPP using a dissolved oxygen change technique (Marzolf *et al*., 1994). Consecutive dawn-dusk-dawn measurements were taken in each mesocosm with a YSI water quality probe (Yellow Springs Instruments, USA). Dissolved oxygen concentration (*μ*mol L^−1^) was then used to calculate net primary productivity and ecosystem respiration, where NPP is the difference in oxygen concentration from the first dawn to dusk measurement divided by elapsed time, and ER is the difference in oxygen concentration from dusk to the second dawn divided by elapsed time. GPP (total photosynthesis) was then calculated as NPP + ER, which assumes that nighttime respiration rates equal those during the day.

Because dissolved oxygen saturation declines with temperature, warming treatments may systematically alter oxygen concentrations not attributed to differences in photosynthesis. We therefore temperature-standardized oxygen concentrations using the Garcia-Benson model (Garcia & Gordon 1992, eqn S3, see Supplementary Methods). In addition, air-water oxygen exchange (reaeration) can alter dissolved oxygen independently of biological fluxes, with the magnitude and direction of exchange depending on the degree of saturation, temperature, and wind speed. At dawn, dissolved oxygen is at its daily minimum following nighttime respiration in the absence of photosynthesis, and is more likely to be undersaturated relative to atmospheric equilibrium; oxygen may therefore diffuse from the air into the mesocosms, causing dawn measurements to slightly overestimate biologically-determined oxygen levels and leading to overestimation of ecosystem respiration. Conversely, daytime photosynthesis drives dissolved oxygen above saturation by dusk, causing outgassing to the atmosphere such that dusk measurements underestimate true photosynthetic oxygen production. Both effects act to underestimate GPP, but are only consequential for our inferences if reaeration varies systematically across the experimental temperature gradient in a way that biases estimated *E*_GPP_ values. We therefore estimated reaeration fluxes for each sampling event (Fig. S3) using an established wind-based gas transfer model (Cole & Caraco, 1998) and quantified their downstream effects on estimated GPP and *E*_GPP_ (see Supplementary Methods and Table S1). As expected, reaeration corrections revealed that dusk dissolved oxygen values were modest underestimates and dawn dissolved oxygen values were modest overestimates, and variation in the magnitude of over- and underestimates was larger among sampling days than among warming treatments (Fig. S3). We confirmed that reaeration rates likely did not bias our GPP and *E*_GPP_ estimates enough to change our main inferences (Table S1; see Supplementary Methods).

### Sampling bacteria, phytoplankton, and zooplankton

Phytoplankton larger than 5*μ*m were the focal group used to test our hypotheses, but we also sampled bacteria and zooplankton to compare the strength and timing of dispersal treatments across trophic levels. To sample bacterial communities, we passed 200 mL of surface water through Sterivex filters (0.22 *μ*m) and stored them in plastic bags at -70°C until DNA extractions for 16S rRNA gene sequencing. We sampled zooplankton communities by passing 10 L of mesocosm water through a 64 *μ*m sieve, returning the water, and preserving trapped zooplankton in 95% EtOH. Methods for bacterial DNA extractions, sequence analysis, zooplankton biomass estimation, and taxonomic identification are described in the Supporting Information.

To estimate phytoplankton biomass, we collected surface samples in 60 mL centrifuge tubes and directly measured chlorophyll-*α* concentration in a Fluoroprobe spectrofluorometer (bbe Moldaenke GmbH, Germany). We checked that variation in chlorophyll-*α* measurements reflected the whole autotroph community by sampling phytoplankton, periphyton, and benthic algae communities, filtering and extracting chlorophyll pigments (Supporting Information), and confirming that their relative biomass contributions were consistent across treatments and over time (Fig. S2). We preserved an additional 60 mL surface water sample from each mesocosm with 1 mL of 25% glutaraldehyde to estimate phytoplankton size spectra and taxonomic composition. We imaged preserved samples at 400x magnification using a FlowCam VS series (Fluid Imaging Technologies, USA). We converted cell volumes (*μ*m^3^) estimated by the Visual Spreadsheet software (Fluid Imaging Technologies, USA) to carbon mass (*μ*g C cell^−1^) by a factor of 0.109 × 10^−6^ (Montagnes *et al*., 1994). We identified and counted phytoplankton sample images in weeks 4, 8, and 12 to the genus or species using the *Key to More Frequently Occurring Freshwater Algae* in Bellinger and Sigee (2015).

To approximate community thermal phenotype composition, we combined the phytoplankton taxonomic count data with published thermal trait data (Thomas *et al*., 2016). The database from Thomas et al. contains thermal response curve parameters from over 6000 growth rate measurements on 240 phytoplankton species, including 11 of the 15 taxa identified in our experiment (Fig. S13, S16). We calculated a Community Thermal Phenotype Index (CTPI) that represents the collection of species’ thermal optima (*T*_opt,*i*_) and thermal widths (*w*_*i*_ = *T*_max,*i*_ − *T*_min,*i*_) weighted by their relative abundance in a given community *c* with *J* total species. Both trait values were standardized across taxa (e.g., 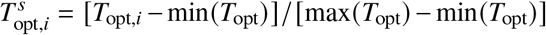), then multiplied to obtain a taxon-specific thermal trait score where the highest value represents the taxon *i* with the highest *T*_opt,*i*_ and widest *w*_*i*_ (eqn 1). Each taxon’s trait score was multiplied by its relative abundance (abundance = *N*_*i,c*_) and scores for all species were summed to obtain raw CTPI:

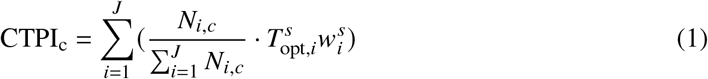

We rescaled eqn 1 as an index where CTPI = 1 captures the community with the highest representation of high *T*_opt_ and wide *w*, and CTPI = 0 has the highest representation of low *T*_opt_ and narrow *w* (eqn S11). We used CTPI to test whether thermal phenotype composition mediates the effect of dispersal on *E*_GPP_ (H2, see below). In order to test the consequences of the four species for which thermal traits were unavailable, we performed a sensitivity analysis to different hypothetical *T*_opt_ values for these species (Fig. S8).

To test for phenotypic sorting into optimal thermal environments, we quantified Community Thermal Match Index (CTMI) which estimates a community’s proximity to its thermal optimum at a given mesocosm temperature given its taxonomic composition. For each community *c*, we weighted the distance of each taxon’s *T*_opt_ from the current temperature *T*_*c*_ by its relative abundance:

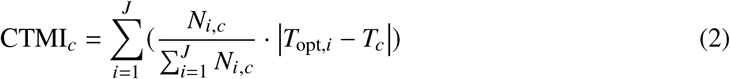

We inverted and rescaled this quantity so that CTMI = 1 represents the community with the best thermal match and CTMI = 0 represents the poorest match across all replicates and time points (eqn S12).

### Analysis

All analyses were performed in R (version 4.4.1, R Core Team 2013). We tested whether different dispersal rates modify the slope describing the temperature sensitivity of GPP, *E*_GPP_ (H1), using a linear mixed effects model (LMM) based on metabolic theory (Allen *et al*. 2005; eqn S2, Supporting Information) with an additional interaction term allowing dispersal to modify the temperature slope:

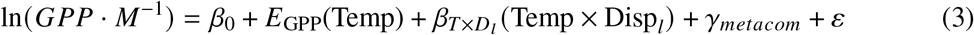

This linearized version of metabolic theory’s canonical model (eqn S2) allows convenient estimation of *E*_GPP_. We used biomass-corrected GPP (*M* = phytoplankton community biomass) to ensure that variation in GPP is attributed to variation in mass-specific photosynthetic rates and not biomass. Temp is standardized Arrhenius temperature 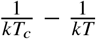, where *T* is mesocosm temperature (Kelvin) and *T*_*c*_ is a standardization temperature 296.15 K. Temp × Disp_*l*_ is an interaction term such that 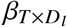 modifies *T*_GPP_ depending on dispersal level *l* (i.e., low or high) relative to a no-dispersal baseline, and *γ*_*metacom*_ is a metacommunity random effect. For our main test of H1, we fit this model separately to data from each week because temporal dynamics violate the model’s steady-state assumptions (Enquist *et al*., 2003) and overwhelm within-time effects of experimental treatments. However, a version of the same model using all time points and a random effect of week (factor) yielded similar results on the effect of dispersal on *E*_GPP_ (Fig. S4).

To test the robustness of inferred time-dependent effects of dispersal on *E*_GPP_, we estimated a version of the model that included data from all weeks with a three-way interaction between dispersal, temperature, and sampling week as a factor; both models yielded the same inferences about H1 (eqn S13, Fig. S4). We also estimated a version of the model with a direct effect of dispersal, but dispersal did not influence the intercept based on our significance criteria outlined below.

Hypothesis 2 implicates several indirect pathways and correlated variables mediating dispersal’s effect on GPP (Directed Acyclic Graph in Fig. S9, Supporting Information). We therefore estimated a piecewise structural equation model (SEM, Grace 2006) to assess the direct and indirect effects of experimental treatments and mediators on GPP across all weeks jointly. We used raw GPP rates (not biomass-corrected) as the response variable given our goal of testing for a mediating effect of biomass. The SEM structure captures the following hypothesized paths, specified as LMMs in Table S2: (1) Temperature and dispersal each indirectly affect GPP via direct effects on phytoplankton biomass, size spectra, and thermal phenotype composition; (2) Zooplankton biomass is directly affected by temperature and dispersal, and in turn indirectly affects GPP through direct effects on phytoplankton biomass, size spectra, and thermal phenotype composition; (3) Nitrogen indirectly affects GPP by increasing phytoplankton biomass, and directly affects GPP by enhancing photosynthetic rates independent of standing biomass (e.g., by overcoming cellular nutrient limitation); (4) Temperature directly affects GPP, which represents its kinetic effect on photosynthesis independent of its effects on community structure. All path coefficients were allowed to vary across weeks by including week as a moderator of all fixed effects (i.e., all predictors were interacted with sampling week as a factor, with week 4 as the baseline), enabling formal tests of whether path slopes changed over the course of the experiment (Table S3). Metacommunity was included as a random effect on intercepts in each submodel. We used CTPI (eqn 1) and size spectra slopes (log size versus log abundance) as proxy variables for thermal phenotype composition and size spectra, respectively. All continuous variables were log-transformed except size spectra slopes and CTPI, and all explanatory variables were scaled.

All LMMs were estimated in a Bayesian framework using the *brms* R package (Bürkner, 2017) with a “cmdstanr” back end. The Bayesian approach helped ensure biologically reasonable parameter estimates and stabilized estimation of slopes and variances when represented by few observations. We sampled from posterior distributions by constructing four chains, each with 5000 steps and a 2000-step warm-up period. Where applicable, we assigned informative priors for the main effects of temperature (mean = 0.65, SD = 0.28), biomass (mean = 0.75, SD = 0.25), zooplankton biomass (mean = -0.5, SD = 0.5), and total nitrogen availability (mean = 0.5, SD = 0.5), and weakly informative priors for phytoplankton community size spectra, thermal phenotype composition, and dispersal (mean = 0, SD = 0.5). We performed a prior sensitivity analysis by comparing posterior estimates with informative priors highlighted above and presented in the Results, wide priors (same mean and twice as large SD), and uninformative priors (mean = 0, SD = 10) for the temperature effect on GPP (Fig. S6).

Model convergence was confirmed by 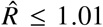 (Gelman–Rubin statistic, Gelman & Rubin 1992). To assess whether explanatory variables had meaningful effects on GPP, we used a practical equivalence test (Kruschke, 2018), which assesses what percentage of posterior distributions fall within the response variable’s region of practical equivalence (ROPE). Paths were considered to have non-zero effects on GPP if < 5% of their posterior 89% highest density interval (HDI) fell within the ROPE (Kruschke, 2018). While a standard recommended ROPE width is ±10% of the response variable’s standard deviation (Kruschke, 2018), we defined a more conservative ROPE of 20% or ±0.20 × SD(GPP) based on a sensitivity analysis (Table S4). Reported values of % of HDI within ROPE refer to a ROPE width of 20% unless otherwise specified. We note that the use of raw GPP in the SEM and mass-corrected GPP in the LMM test of H1 reflect separate inferential goals; metabolic theory temperature predictions about *E*_GPP_ (eV) are defined for per-unit-biomass productivity, but our ability to distinguish effects of multiple mediators of dispersal on GPP using SEM requires biomass to be separated from the GPP response variable.

As additional tests of dispersal’s homogenizing effect on community structure, we calculated the coefficient of variation (CV) for biomass, size spectra slopes, and CTPI within each metacommunity, then tested for differences among dispersal treatments with 1-way ANOVA and Tukey’s Honestly Significant Difference (HSD) post-hoc test. To compare homogenizing effects on taxonomic composition across trophic levels (bacteria, phytoplankton, zooplankton), we performed Permutational Analysis of Multivariate Dispersion (PERMDISP) on taxonomic count data to compare mean within-metacommunity dispersion (community distance to metacommunity centroid) among dispersal treatments with 1-way ANOVA and Tukey’s HSD. To test whether higher dispersal weakens temperature effects on size spectra slopes, we estimated LMMs of log abundance versus log size with size-by-temperature and size-by-dispersal interactions (eqn S14, Table S7).

## Results

Changes in communities and GPP reflected successional dynamics (monotonic biomass accumulation, species turnover) and responses to thermal fluctuations over the 12-week experiment. Phytoplankton biomass and GPP rates increased steadily in all mesocosms from weeks 2 to 8 (Fig. 2). From week 8 onward, mean GPP rates either remained stable or decreased as ambient temperatures declined, despite continued increases in biomass. Chlorophytes consistently dominated autotroph community biomass, though diatoms, cryptophytes, chrysophytes, and cyanobacteria were also present (Fig. S7).

**Figure 2.**
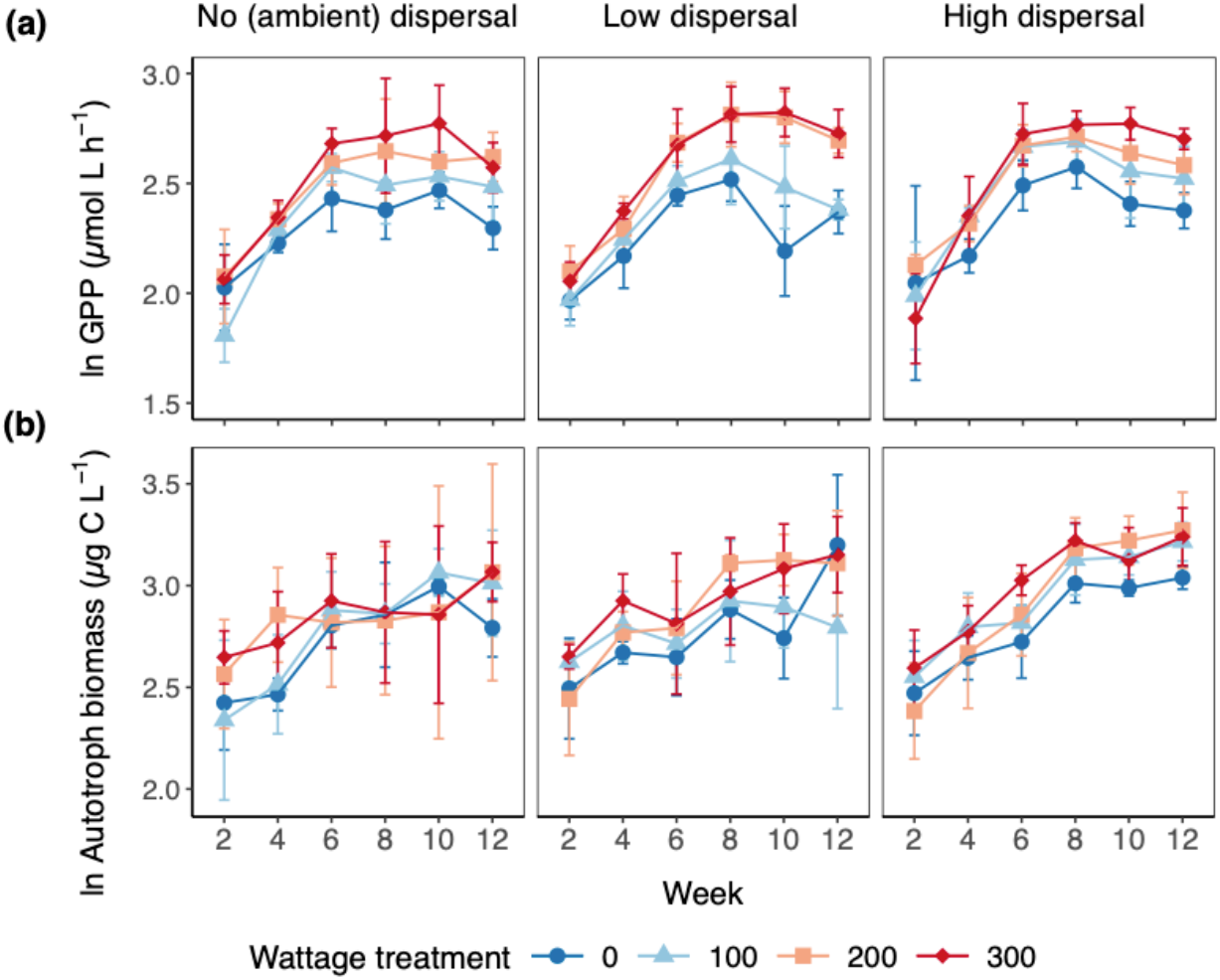
Changes in **(a)** gross primary productivity (mean ±SD) and **(b)** autotroph biomass (mean ±SD) by dispersal treatment (panel columns) over the 12-week experiment. Point shapes and colours indicate warming treatment.

### H1: Dispersal modifies the thermal sensitivity of GPP

We found that *E*_GPP_ differed among dispersal treatments only briefly after the unplanned heatwave (Fig. 3). In week 8, *E*_GPP_ was significantly weaker in high-dispersal treatments compared to other dispersal treatments (posterior mean *E*_GPP_ = 0.17 [89% HDI -0.29, 0.70], 0% HDI within ROPE). The reduced *E*_GPP_ under high dispersal in week 8 remained consistent in direction, magnitude, and statistical support regardless of prior specification (i.e., strong, weakly informative, or uninformative; Figure S6). The thermal sensitivity of mass-specific GPP changed over time regardless of dispersal, such that *E*_GPP_ was significantly positive from week 6 onwards, peaked in week 10 (0.72 [0.44, 0.99], 0% within ROPE) then declined by week 12 (0.43 [0.14, 0.72], 0% within ROPE). Mass-specific GPP was not significantly temperature-dependent in weeks 2 (0.33 [-0.03, 0.71], 7.21% within ROPE) or 4 (0.20 [-0.03, 0.45], 9.41% within ROPE); this may reflect a lower signal-to-noise ratio in chlorophyll measurements due to low biomass early in the experiment (Fig. 2b) or delayed responses to temperature treatments. This effect was also supported in a version of the model estimated with data from all weeks (Fig. S4) and a model using raw GPP instead of biomass-corrected GPP as the response variable (Fig. S5).

**Figure 3.**
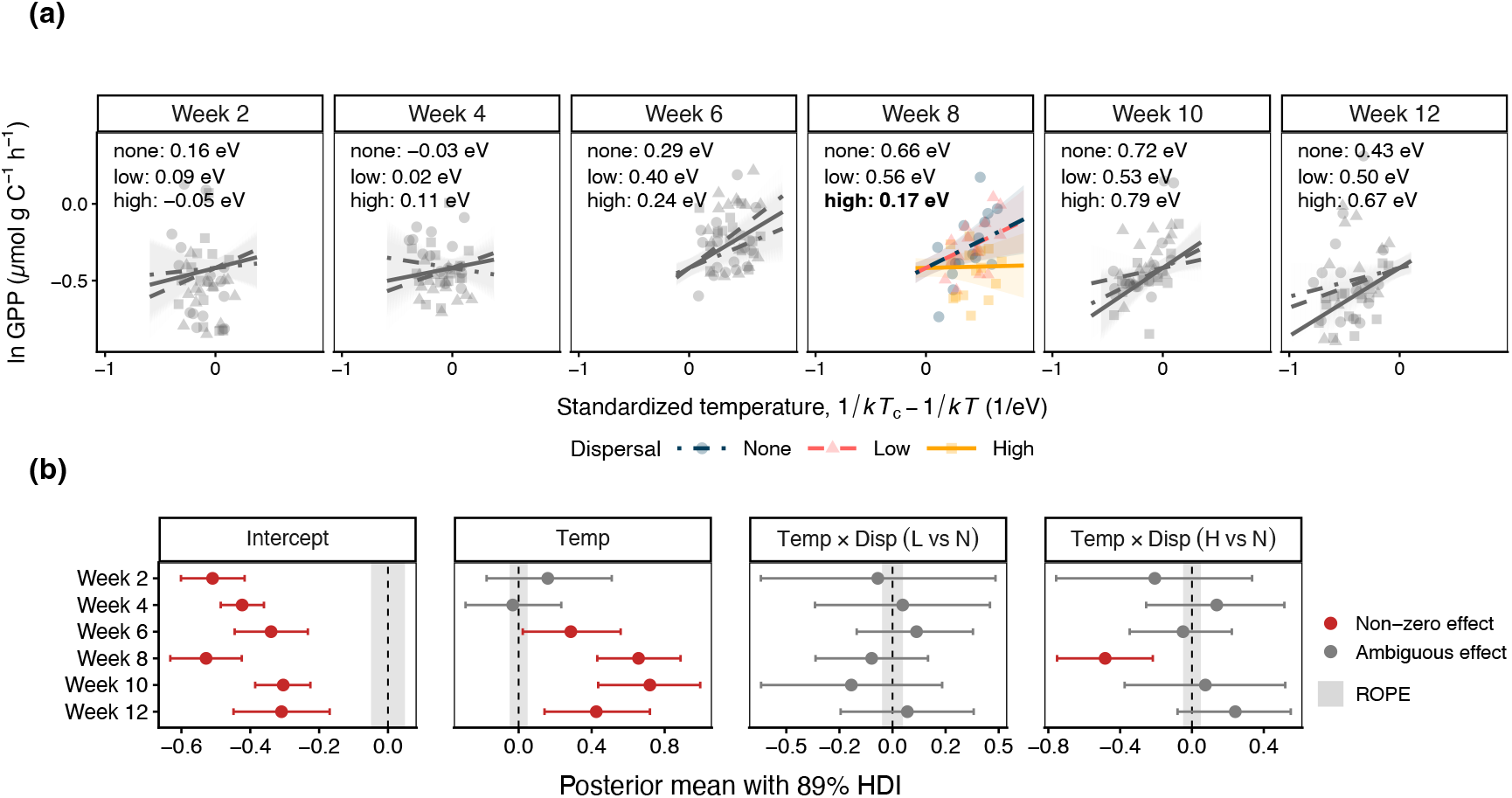
Thermal sensitivity of biomass-specific gross primary productivity by dispersal treatment and sampling week. **(a)** Coloured points and lines indicate weeks where dispersal modified the slope (*E*_GPP_) of the temperature-GPP relationship. Grey lines indicate weeks where *E*_GPP_ did not significantly differ among dispersal treatments. **(b)** Forest plot of posterior estimates ( ±89% HDI) for each fixed effect in the LMMs represented in panel (a). Grey shaded regions represent Region of Practical Equivalence. The “Temp × Disp” panels represent pairwise differences in the slope for temperature versus GPP (i.e., *E*_GPP_) among the three dispersal treatments.

### H2: Dispersal affects GPP via biomass, size spectra, and thermal phenotypes

There was a strongly supported (credibly non-zero) direct effect of temperature on GPP across all weeks (0% of HDI within ROPE, Table 1, Fig. S9). This strong direct effect likely obscured clear support for an indirect effect of temperature on GPP via mediators, despite strongly supported temperature effects on phytoplankton biomass in weeks 4 and 12, phytoplankton thermal phenotype composition in week 8, and zooplankton biomass in all three weeks (< 5% of HDI within ROPE, Table S3). Temperature effects on phytoplankton thermal trait composition in weeks 4 and 12, biomass in week 8, and size spectra in all three weeks were sensitive to ROPE width and therefore ambiguous; effects were supported at more lenient ROPE widths of 5% or 10% of SD(GPP), but were indistinguishable from zero at the stricter ROPE width of 20% (Table S4). It is possible that the modest effective sample size for testing dispersal effects may account for the wide credible intervals on several SEM paths, such that our analysis failed to detect some non-zero indirect path effects.

**Table 1:** Direct and indirect paths affecting GPP by week based on SEM analysis. Bold rows represent paths with non-zero effects on GPP (< 5% HDI within ROPE = ±0.20×SD(GPP)).

| Path | Week | Posterior mean | 89% HDI | % HDI in ROPE |
| --- | --- | --- | --- | --- |
| High dispersal → GPP (indirect) | 4 | 0.022 | [-0.011, 0.061] | 77.7 |
|  | **8** | **0.121** | **[0.053, 0.198]** | **0.0** |
|  | 12 | 0.034 | [-0.014, 0.088] | 58.6 |
| Low dispersal → GPP (indirect) | 4 | 0.015 | [-0.021, 0.058] | 86.5 |
|  | 8 | 0.063 | [-0.002, 0.131] | 34.2 |
|  | 12 | 0.009 | [-0.040, 0.061] | 83.4 |
| **Temp → GPP (direct)** | **4** | **0.309** | **[0.152, 0.470]** | **0.0** |
|  | **8** | **0.579** | **[0.413, 0.743]** | **0.0** |
|  | **12** | **0.665** | **[0.514, 0.813]** | **0.0** |
| Temp → GPP (indirect) | 4 | 0.063 | [-0.005, 0.146] | 34.6 |
|  | 8 | 0.085 | [-0.045, 0.219] | 31.9 |
|  | 12 | 0.032 | [-0.067, 0.137] | 41.3 |

High dispersal had a strongly supported indirect effect on GPP in week 8 (0% HDI within ROPE), most likely through elevated phytoplankton biomass (Fig. S9b, Table S3). The effect of high dispersal on thermal trait composition was sensitive to ROPE width in week 8, such that the contribution of thermal traits to the indirect effect on GPP was ambiguous. There was no support for an indirect effect of dispersal on GPP in weeks 4 or 12 at any ROPE width (Table S4). Nitrogen concentration did not affect GPP at any time, nor was there support for zooplankton affecting any aspect of the phytoplankton community (Fig. S9, Table S3).

#### Biomass

Higher dispersal rates significantly reduced within-metacommunity biomass variance (CV) in weeks 8 through 12 (Fig. 4, Table S5). High-dispersal communities also had significantly lower variance in biomass and GPP among warming treatments in week 8, whereas lower dispersal treatments had consistently larger variances (Fig. 2, Levene’s test of Equality of Variances, Table S6). Within-metacommunity biomass variance (CV) and within-metacommunity *E*_GPP_ (not biomass-corrected) were positively correlated in week 8 (Fig. S10), suggesting that this variance-reducing effect of high dispersal may have contributed to the reduced *E*_GPP_.

**Figure 4.**
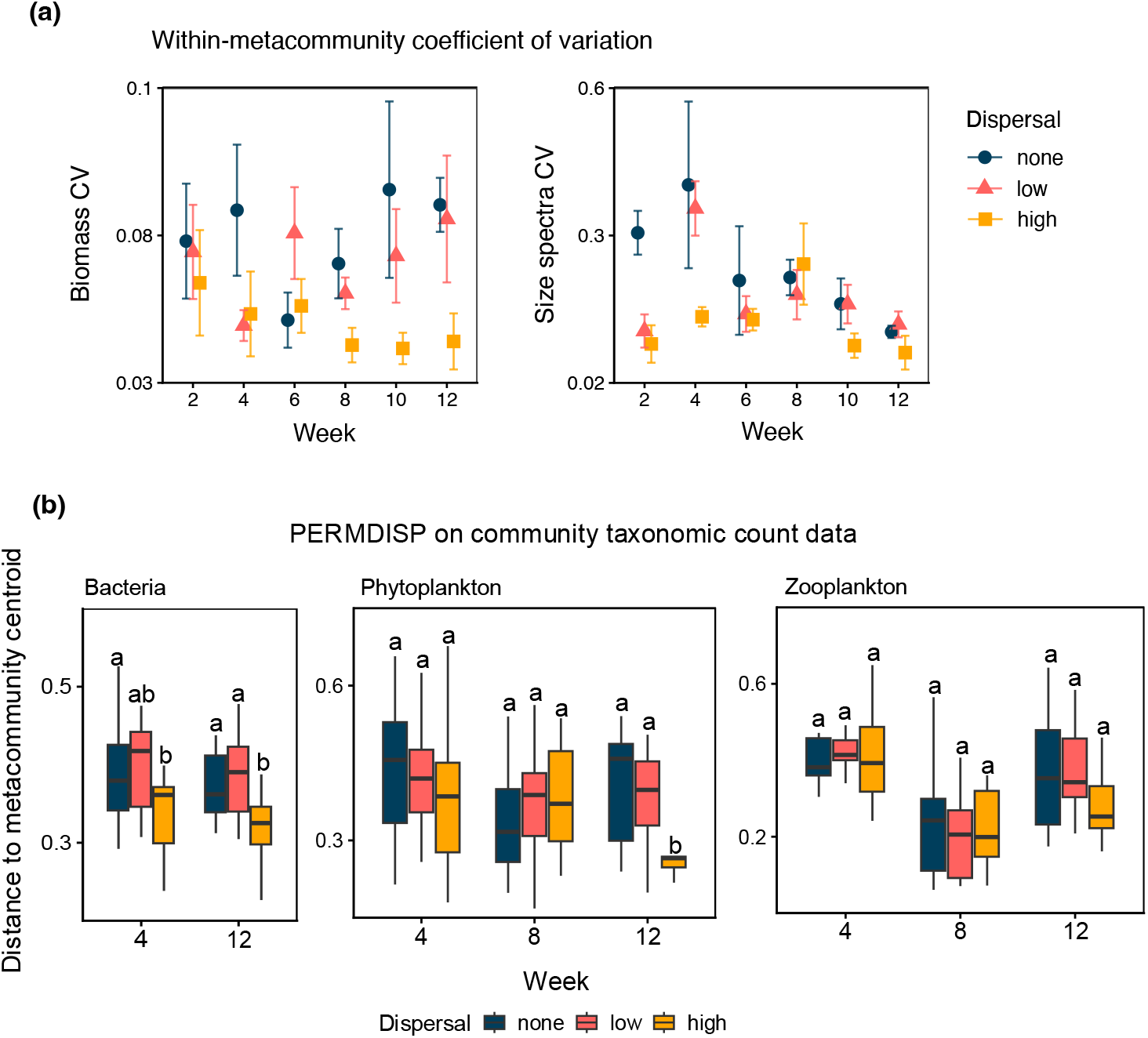
Within-metacommunity variation in biomass, size spectra, and taxonomic composition. **(a)** Within-metacommunity coefficient of variation (CV ± SE) for phytoplankton community biomass and size spectra slopes. Each point represents the CV within a single metacommunity linked by dispersal, such that the range of temperatures (warming treatments) is equal among dispersal groups. **(b)** Within-metacommunity multivariate dispersion (PERMDISP) on taxonomic composition of bacteria, phytoplankton, and zooplankton. Points reflect the distance between a community and the centroid of its respective metacommunity. Letters indicate dispersal treatment differences based on Tukey HSD post-hoc test on 1-way ANOVA.

#### Size spectra

Phytoplankton communities were consistently comprised of many small cells and few large cells (negative size spectra slope, right-skewed distribution, Fig. S11). Contrary to our predictions, temperature did not affect size spectra (LMM: size x temp, 75–100% of HDI within ROPE, Table S7), nor did it interact with dispersal to affect size spectra (LMM: size x temp x disp, 69–100% of HDI within ROPE, Table S7). Dispersal, however, modified size spectra independently of temperature; low dispersal yielded significantly shallower size spectra slopes and high dispersal yielded steeper ones relative to no dispersal in a subset of weeks (Fig. S12, Table S7). High dispersal also reduced within-metacommunity size spectra variance, though statistical support for this was only detected in week 2 (Tukey HSD, df = 12, p = 0.01, Fig. 4, Table S5).

Though temperature did not affect size spectra, temperature affected other aspects of community size structure. Average cell volumes were smaller at the warmest temperatures within weeks 8 and 10, reflecting a temperature-size rule scenario (Fig. S12b). Temperature had an opposite effect on average cell size over time, with the largest individual cell volumes observed during the warmest period (week 8) and smallest during the coolest period (week 12) (Fig. S12b). Zooplankton biomass followed a similar trend such that phytoplankton cell size was positively correlated with zooplankton biomass over time but not within weeks (Fig. S12c).

#### Thermal phenotype composition

We tested the prediction that thermal phenotype composition best matches local temperatures under moderate dispersal, and that higher dispersal weakens whichever thermal sensitivity pattern (hotter-is-better versus compensation) occurs under moderate dispersal. Thermal optima for phytoplankton taxa in the mesocosms ranged from 16.9–30.8 °C based on published thermal response curve assays (Thomas *et al*., 2016). Maximum growth rates increased with thermal optima on a log-linear scale (Fig. S13); this is expected to facilitate a hotter-is-better *E*_GPP_ (Fig. 1c) if species’ occurrences reflect phenotypic sorting into optimal thermal environments (here temperature treatments). We did not observe strong differences in thermal optima representation (i.e., relative abundances) among warming treatments in weeks 4 and 12. In week 8, low-dispersal communities showed the greatest temperature-driven differences in thermal optima representation, with the highest abundances of more heat-tolerant species (*T*_*opt*_ ≈ 30 °C) in the 300 W communities and highest abundances of colder-adapted species (*T*_*opt*_ < 20 °C) in the 0 W communities (Fig. 5a). High-dispersal communities had the smallest differences in thermal optima representation among warming treatments in week 8 (overlapping distributions in Fig. 5a), possibly due to homogenization.

**Figure 5.**
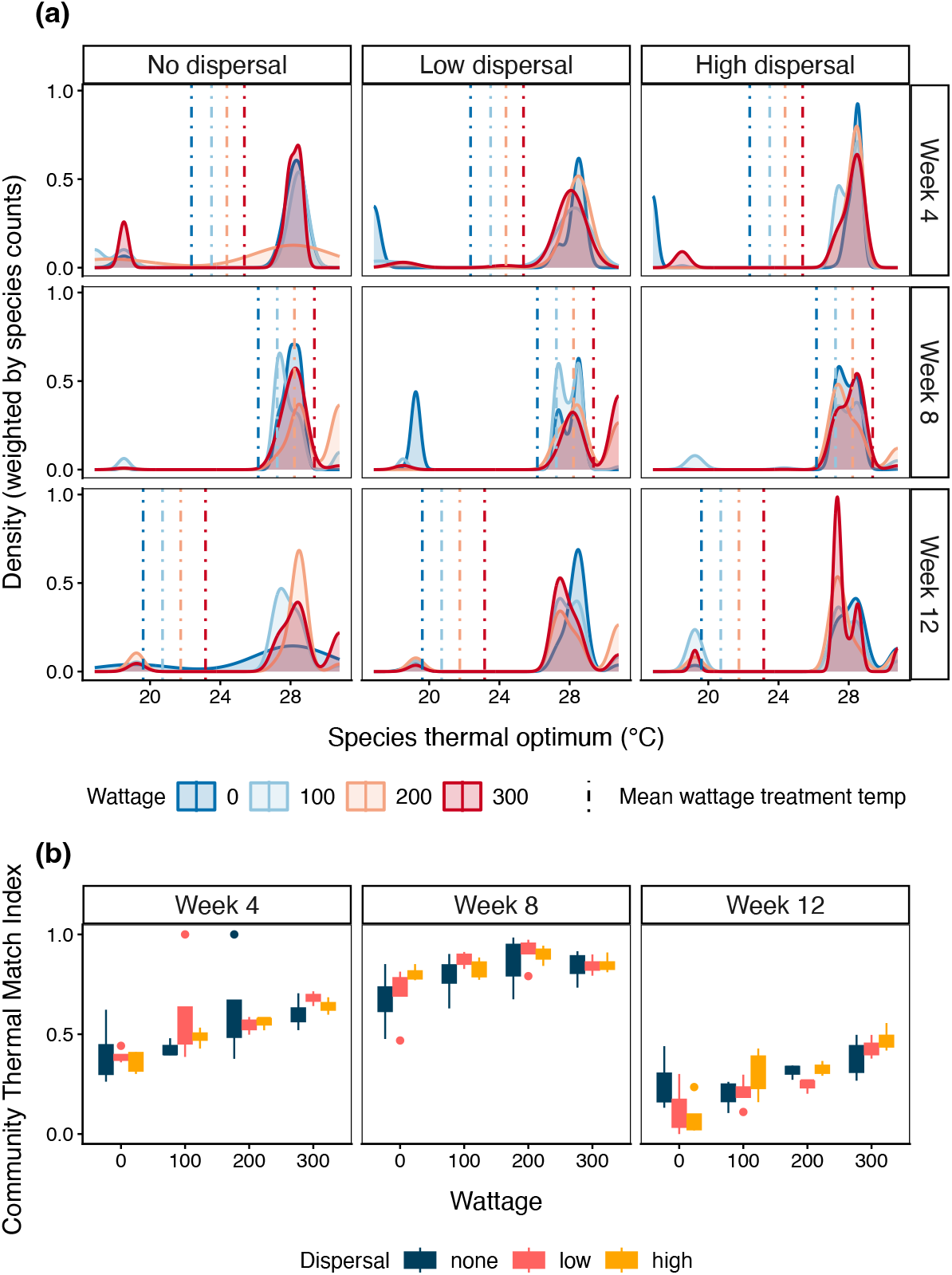
Summaries of thermal phenotype distributions and community-level thermal match across experimental treatments and weeks. **(a)** Distribution of thermal optima within communities at different warming treatments based on relative abundances of phytoplankton taxa. Vertical dashed lines represent the average mesocosm temperature of a given warming (wattage) treatment that day. **(b)** Mean community thermal phenotype match index (CTMI) by wattage and dispersal treatment. Values of 1 represent the best thermally matched community across all treatments and times, and 0 represents the worst thermally matched community based on the relative abundances of species and their thermal optima.

Communities were overall closest to their thermal optima in week 8 (Fig. 5b). Communities in 200 W treatments had the highest thermal match, and the relatively lower thermal match in the 300 W treatments suggests those temperatures were beyond species’ thermal optima. In other weeks, the 300 W communities had the highest community thermal match.

Within-metacommunity variance in CTMI did not statistically differ with dispersal (Tukey HSD, df = 12, p > 0.05, Table S5), however, high dispersal reduced within-metacommunity variance in taxonomic composition (Fig. 4b). The strength and timing of this effect were weaker and delayed with increasing trophic level. In the bacterial communities, high dispersal significantly reduced within-metacommunity multivariate dispersion in taxonomic composition compared to low (Tukey HSD on PERMDISP, df = 46, p = 0.001) and no dispersal (Tukey HSD on PERMDISP, df = 46, p = 0.008; see Fig. S14 for bacterial community composition). Among phytoplankton communities, this homogenizing effect was only detected in week 12, where dispersion within high-dispersal metacommunities was significantly reduced compared to no dispersal and low dispersal ones (Tukey HSD on PERMDISP, df = 44, p < 0.05). Zooplankton in high-dispersal metacommunities showed marginally lower dispersion in week 12 (Tukey HSD on PERMDISP, df = 45, p = 0.06), but not in the weeks prior.

## Discussion

Metabolic scaling theory posits that the thermal sensitivity of individual metabolism determines how ecosystem productivity responds to warming across broad spatiotemporal extents (Allen *et al*., 2005; Enquist *et al*., 2003; Yvon-Durocher *et al*., 2012). Ecological and evolutionary dynamics can alter these scaling relationships through plasticity (Kordas *et al*., 2022a), adaptation (Kontopoulos *et al*., 2020; Padfield *et al*., 2016; Schaum *et al*., 2017), resource dynamics (Anderson-Teixeira *et al*., 2008; Vinton & Vasseur, 2022; Yvon-Durocher *et al*., 2012), and species interactions (Cruz *et al*., 2023; Garcia *et al*., 2023), but whether dispersal, a similarly widespread and important process, modifies the thermal sensitivity of ecosystem productivity has received less attention. To our knowledge, this study provides the first explicit test.

We found that warming treatments had the strongest and most consistent direct effect on GPP (Fig. S9), reflecting temperature’s instantaneous kinetic effects on photosynthetic rates (Arrhenius, 1889; Crozier, 1926; Gillooly *et al*., 2001; Hutchins, 1947). High dispersal temporarily weakened the thermal sensitivity of GPP (*E*_GPP_, slope of the GPP-temperature relationship) after a heatwave as a result of lower mass-specific productivity at the warmest temperatures relative to lower dispersal magnitudes (Fig. 3). Differences in GPP and community structure among dispersal treatments were consistent with a regional homogenizing effect and *E*_GPP_ reflecting a hotter-is-better scenario (Fig. 1). Aside from this temporary response to an extreme thermal fluctuation, and consistent with metabolic theory’s steady-state assumptions, the thermal sensitivity of GPP was robust to dispersal.

Dispersal can buffer communities against extirpation after disturbance, but spatial insurance effects on abundance and biomass may not necessarily translate to enhanced productivity, particularly after heat stress (de Boer *et al*., 2014; Eggers *et al*., 2012; Thompson & Shurin, 2012). Another metacommunity experiment with microalgae showed that manipulating dispersal after a heatwave spread a heat-tolerant but low-productivity species, depressing biomass (Eggers *et al*., 2012). Our results differed in that high dispersal slightly elevated biomass (Fig. S9), but we similarly found that mass-specific primary productivity was depressed post-heatwave (Fig. 3). High dispersal significantly increased the relative abundance of *Oocystis* sp. in week 8, based on statistical support from an LMM (Fig. S15; see Supporting Information). *Oocystis* has a large cell size and wide thermal range but lower *T*_opt_ and maximum growth rate than other taxa in the experiment (Fig. S13). High dispersal may therefore have facilitated higher standing biomass through the spread of larger phytoplankton that have a lower propensity for ambient dispersal, but in so doing spread poorly adapted thermal phenotypes into the warmest mesocosms. Support for this hypothesis is ambiguous based on our results; the effect of high dispersal on thermal phenotype composition was sensitive to ROPE width in week 8, and this did not translate into a statistically supported indirect effect of dispersal on GPP via thermal phenotypes (Table 1). This may be due to the limited sample size of each temperature-by-dispersal treatment combination, or inaccuracies in the CTPI metric itself. While our sensitivity analysis of CTPI estimates to the four missing species traits confirmed that the rank-order of CTPI among replicates is broadly robust to different hypothetical *T*_opt_ values for these species, it also showed that the highest CTPI values used in our SEM may be overestimates if all missing species had thermal optima of 16°C (Fig. S8). This value is low for temperate freshwater phytoplankton (Thomas *et al*., 2016) and thus unlikely to be the true *T*_opt_ of all four missing species. We nevertheless acknowledge that uncertainty in CTPI estimates along with the statistically ambiguous effect of dispersal in the SEM analysis suggest that the hypothesized mechanism remains speculative given the data.

High dispersal reduced differences in thermal optima representation among warming treatments and reduced community thermal match compared to low-dispersal treatments (Fig. 5). We thus hypothesize that a hotter-is-better *E*_GPP_ pattern was facilitated by lower dispersal rates and temporarily weakened by high dispersal spreading less heat-tolerant phenotypes into the warmest (300 W) mesocosms, lowering mass-specific productivity (Fig. 3). After weeks of steady cooling, *E*_GPP_ under high dispersal was indistinguishable from that of other dispersal treatments. In experimental diatom metacommunities subjected to a heatwave, de Boer *et al*. (2014) found that dispersal reduced equilibrium biomass under stable conditions, but enhanced post-disturbance biomass recovery. If this mechanism occurred in our experiment, then the negative effect of high dispersal on productivity in week 8 through the spread of cold-adapted phenotypes in the 300 W communities could have facilitated its recovery in week 10. Alternatively, dispersal’s transient effect on *E*_GPP_ could be an exceptional case that only applies when temperatures exceed species’ thermal optima and poorly adapted thermal phenotypes are dispersed throughout the metacommunity (Fig. 5b), but under benign conditions dispersal’s homogenizing effect does not significantly affect productivity. We stress that the inferred transient effect of high dispersal on *E*_GPP_ reflects the specific successional dynamics of our experimental communities. Given the heatwave was not experimentally planned or replicated, the generality of our result showing that high dispersal weakens *E*_GPP_ after a heatwave cannot be confirmed. In theory, the hypothesized mechanism in the present study may extend to other perturbations where dispersal may act on variation of relevant traits (i.e., cold tolerance traits under a cold spell, nutrient acquisition traits under a nutrient pulse). Future work that explicitly estimates changes in trait composition and exposes communities to replicated perturbations may clarify the generality of our results.

The SEM did not support effects of zooplankton on phytoplankton size, which may reflect the limited sampling effort of the zooplankton community (weeks 4, 8, and 12 only). Trends over time suggested possible temperature-mediated top-down control of phytoplankton community size structure as found in multi-trophic warming experiments (Padfield *et al*., 2018; Yvon-Durocher *et al*., 2015). Higher zooplankton abundances combined with intensified grazing rates in week 8 may have caused this pattern. In another study focused on zooplankton communities in this experiment, we report temperature-driven differences in zooplankton community composition within and across time; *Daphnia* sp. and calanoid copepods peaked in abundance at cooler temperatures, whereas smaller cladoceran *Diaphanosoma* sp. dominated at the warmest temperatures in week 8 (Thompson *et al*., 2024). Here, higher mean cell sizes were driven largely by higher abundances of colonial green alga *Crucigenia* sp., possibly reflecting selection for larger cells and coloniality under heightened grazing pressure. The divergent relationships between temperature and phytoplankton size within versus across time may suggest that trophic effects occur on a longer time scale than do physiological temperature effects.

Our results suggest size-dependent sensitivity to dispersal; high dispersal steepened phytoplankton size spectra in several weeks (Fig. S12a) and had systematically weaker effects on taxonomic composition with increasing trophic level (Fig. 4b). Body size and generation time are the main traits underlying this pattern. Unicellular organisms have high population abundances and short generation times resulting in higher colonization success and quickly detectable changes in abundance and diversity (Finlay, 2002; Thompson & Gonzalez, 2017). Even within microbes (i.e., bacteria versus phytoplankton), dispersal ability varies significantly among taxa with different life history and morphological traits (Echenique-Subiabre *et al*., 2025). Zooplankton have lower population abundances and longer generations, such that experimental dispersal may have weaker, slower, or undetectable effects on community composition, even under environmental stress (Limberger *et al*., 2019). While our study examined dispersal effects on productivity, this result suggests future directions investigating whether dispersal has different effects on community respiration or metabolic fluxes involving other trophic levels with different dispersal sensitivities.

We acknowledge several data availability limitations arising from the time and resource constraints inherent to a large-scale outdoor mesocosm experiment. The relatively small number of metacommunity replicates per distinct dispersal-by-temperature treatment combination (n = 4) may have contributed to several ambiguous effect paths in our SEM analyses. Our use of a Bayesian framework aimed to mitigate the impact of limited replication, and our distinction between clear and ambiguous path effects was informed by several sensitivity analyses (Table S4, Fig. S6). We also note that dispersal of dissolved compounds such as dissolved organic carbon, allelopathic substances, or other metabolites may also have contributed to differences in mass-specific productivity among dispersal treatments, though we did not have the data to test this. A limitation in our dawn-dusk-dawn method of estimating GPP arises from potential systematic differences in reaeration rates at different times of day when oxygen levels tend to be over- or undersaturated. While this could have caused greater underestimation of GPP in warmer tanks, a re-analysis of the LMM testing H1 with reaeration-corrected GPP estimates yielded the same results about effects of dispersal on *E*_GPP_ (Table S1). Finally, our use of published thermal traits measured in laboratory conditions may not necessarily reflect realized thermal traits in communities, which may change via plasticity, adaptation, and biotic interactions (Garcia *et al*., 2022; Kremer *et al*., 2018; Schaum *et al*., 2018). Despite this, previous studies show that laboratory-derived trait estimates are surprisingly effective at explaining phytoplankton distributions (Thomas *et al*., 2012) and seasonal community dynamics (Edwards *et al*., 2013a,b) in nature. This body of work suggests that capturing broad trait variation patterns suffices to explain interspecific differences in growth and abundance, and exact quantitative trait values may not be necessary.

## Conclusion

Our findings suggest that metabolic theory’s temperature scaling predictions are robust under steady-state conditions, but may depend on dispersal magnitude after an extreme thermal fluctuation. Empiricists applying metabolic scaling predictions in natural systems should therefore situate their observations in the context of recent perturbations, successional dynamics, and dispersal limitation. Our observation that no dispersal communities showed higher variance in community structure and productivity suggests that temperature effects may be less predictable in dispersal-limited systems, with implications for systems facing the dual pressures of habitat fragmentation and changing thermal regimes. Continued synthesis of metabolic theory with community ecology will yield improved insights into ecosystem responses to short-term environmental change at local and regional scales.

## Acknowledgments

We are grateful to Felipe Amadeo for identifying zooplankton taxa. We also thank Bianca Trevizan Segovia, Claire Atkinson, Katie Tjaden-McClement, Kendrix Murray, and Natalie Westwood for assistance during the experiment. KS was supported by an NSERC Alexander Graham Bell Canada Graduate Scholarship. PLT was supported by Killam and NSERC postdoctoral fellowships. JRB was supported by a Nereus Fellowship. The experiment was funded by an NSERC Discovery Grant (2018-05043) to MIO.

## Statement of Authorship

JRB, MIO, and KS conceived of the study. PLT and MIO conceived of and designed the experiment with input from CF and JRB. MIO provided funding. PLT, CF, JRB, and KS performed field collections. PLT, CF, JRB, KED, KS, and EY ran the experiment. EY and ASJ provided 16S sample preparation and bioinformatics, with supervision from LWP. MH performed flow cytometry. KED provided nutrient analyses. KS performed phytoplankton identification and data analysis, generated figures, and wrote the manuscript. All authors provided manuscript edits.

## Data Accessibility Statement

Data and code to reproduce analyses are available from https://github.com/keilastark/dispersal_temp_scaling_experiment and will be published to a permanent repository after publication.

