## Supporting Information for "Dispersal transiently modifies the temperature dependence of ecosystem productivity after an extreme thermal fluctuation"

### Supplementary Methods

#### *Modelling the metabolic scaling of GPP*

At temperatures below an individual's thermal optimum, whole-organism photosynthetic rate $Ps_i$ depends on mass and temperature according to:

$$Ps_i = Ps_i(T_c)m^{\alpha}e^{E_{Ps}(\frac{1}{kT_c}-\frac{1}{kT})} \tag{S1}$$

where $Ps_i(T_c)$ describes metabolic rate at an arbitrary normalization temperature, $T_c$, $m$ is body or cell mass, $\alpha$ is an allometric scaling exponent, $T$ is temperature in Kelvin, $k$ is the Boltzmann constant, and $E_{Ps}$ (eV) is the thermal sensitivity parameter for photosynthesis, which describes the temperature dependence of $Ps_i$, and has been historically referred to as an activation energy due to its origins in enzyme kinetics (Gillooly *et al.*, 2001).

The mass and temperature dependence of gross primary productivity (GPP) is modelled using the mass-specific temperature dependence of individual metabolic rates. To apply this to aquatic systems, we assume that the community-level metabolic rate is approximated by the biomass-weighted sum of all constituent populations or size classes per unit volume (V):

$$\text{GPP} = \frac{1}{V}\sum_{i=1}^{n} Ps_i(T_c)M_{\text{comm}}\langle m_i^{\alpha_{Ps}-1}\rangle e^{E_{\text{GPP}}(\frac{1}{kT_c}-\frac{1}{kT})} \tag{S2}$$

where $T_c$ is a normalization temperature (i.e., temperature at which the intercept occurs), $M_{\mathrm{comm}}$ is the total biomass of the autotroph community, and $m_i$ is the mean individual mass of species $i$. Thermal sensitivity parameter $E_{\mathrm{GPP}}$ describes the temperature dependence of GPP. Its value is often assumed to equal that for individual photosynthesis ($E_{\mathrm{GPP}} = E_{Ps}$; Allen *et al.* 2005; Gillooly *et al.* 2001). There is evidence for this in macroecological studies (Enquist *et al.*, 2009), but empirical evidence collected on shorter spatial and temporal scales suggests that $E_{\mathrm{GPP}}$ can change and take on more variable values due to adaptation, plasticity, and enhanced resource availability (Cross *et al.*, 2015; Padfield *et al.*, 2017; Welter *et al.*, 2015). All thermal sensitivity values reported in our study reflect $E_{\mathrm{GPP}}$ only, as we did not isolate and measure photosynthetic rates in individual cells.

The term $M_{\mathrm{comm}}\langle m_i^{\alpha_{Ps}-1}\rangle$ represents size-corrected biomass, which facilitates even comparisons of metabolic capacity among communities with similar biomass but different size structures (Barneche *et al.*, 2014; Padfield *et al.*, 2018; Yvon-Durocher & Allen, 2012). Empirical estimates for the mass-dependence ($\alpha$) of photosynthetic rate range between 0.75-1 (Finkel *et al.*, 2010; Marañón, 2015; Padfield *et al.*, 2018); in cases where $\alpha$=1, a size-correction is not necessary since $m_i^{\alpha_i-1}$ becomes 1, and GPP is simply a function of total community biomass $M_{\mathrm{comm}}$. We did not size-correct community biomass in our analyses because doing so would have confounded size-mediated dispersal effects on GPP. The biomass-correction for GPP in the test of H1 (eqn 3 in main text) is reasonable given empirical support for $\alpha$=1 in eukaryotic phytoplankton (Finkel *et al.*, 2010).

#### *Bacterial 16S sequencing and bioinformatics*

In preparation for DNA extraction, Sterivex filters were opened in sterile conditions following Cruaud et al. (2017). Half of each filter was placed in a 96-well extraction plate while the other half was saved and stored at -70°C as a backup. We extracted DNA with the Qiagen PowerSoil® HTP 96 Well DNA and Qiagen PowerSoil Pro® Extraction Kits following the manufacturer's protocol. Samples were randomly sorted on two extraction plates. We amplified the V4 region of the 16S rRNA gene and ran Phusion Flash PCR (30 cycles) with each reaction containing 10 mg $mL^{-1}$ BSA, 6 $\mu$L water, 10 mM 1 $\mu$L forward primer (515F: GTGYCAGCMGCCGCGG TAA with Illumina adapters and a 12nt barcode), 10mM 1 $\mu$L reverse primer (806R: GGACT ACHVGGGTWTCTAAT), and 1 $\mu$L DNA. Amplified DNA was imaged on a 1% agarose gel (2.5 g agarose + 250 mL TAE) and cleaned. Samples extracted with the Powersoil HTP 99 well kit were purified using the Qiagen Ultra Clean 96 PCR kit, and samples extracted with Qiagen PowerSoil Pro were purified using QIAquick PCR purification kit. Purified DNA amplicons were quantified using PicoGreen and pooled to equal concentration (17 ng). The final pool was submitted for Illumina MiSeq amplicon sequencing to the Integrated Microbiome Resource facility at the Centre for Comparative Genomics and Evolutionary Bioinformatics at Dalhousie University (Halifax, Canada) according to standard protocols (Comeau *et al.*, 2017).

Raw sequencing reads from bacterial communities were demultiplexed using the idemp workflow (Wu, 2014), then processed into amplicon sequence variants (ASVs) using the DADA2 16S pipeline, including trimming to a minimum sequence length of 150 bp and removing chimeras. ASVs were assigned taxonomy using the SILVAv132 (Callahan, 2018) database clustered at 97% similarity. ASVs with fewer than 100 reads in the dataset were filtered out, as were ASVs unassigned at the domain level. Following this, data were loaded into phyloseq (version 22 3.6.3, McMurdie & Holmes 2013) and chloroplasts and mitochondria were removed. For diversity analyses, samples were rarefied to 20,000 reads, and samples with fewer than 20,000 reads were removed from the analysis, including all negative controls (5 extraction negatives and 2 PCR negatives). Bacterial ASV identities are described in detail in Yangel 2021.

#### *Zooplankton identification and biomass quantification*

To estimate zooplankton community biomass, preserved samples were split using a Folsom Plankton Splitter (Aquatic Research Instruments, USA) then counted until a total abundance of 100 individuals per species was reached; this number was divided by the proportion of sub-samples counted to estimate total abundance. The remaining sub-samples were counted for all species that did not reach this 100 individual threshold. Samples were dried in a 60°C oven for 48 hours and weighed to estimate zooplankton biomass density ($\mu$g $L^{-1}$). Zooplankton were identified under a stereo microscope (Leica M165C) at 10x magnification to the lowest taxonomic group possible. The zooplankton community consisted of 10 broad taxonomic groups including cladocerans (*Daphnia* sp., *Diaphanosoma* sp., *Scapholeberis* sp., *Simocephalus* sp.), ostracods, copepods (orders cyclopoida and calanoida), water mites, and midge (Chaoboridae and Chironomidae) larvae (Thompson *et al.*, 2024).

#### *Comparing autotroph biomass in different parts of the mesocosm*

Phytoplankton communities were sampled by submerging 60 mL centrifuge tubes by hand into the middle of each mesocosm. Periphyton communities were sampled from 20 x 20 cm plastic square tiles subdivided into equal rectangular strips and hung with fishing line along mesocosm walls. At each sampling time we scraped one strip with a toothbrush and collected dislodged periphyton in a centrifuge tube. Benthic microalgal communities were sampled from 20 x 20 cm ceramic tiles subdivided into rectangular strips placed on the mesocosm floor using the same scraping method. Samples were passed through a GF/F glass filter paper (0.7 $\mu$m pore size) before extracting chlorophyll pigments in acetone for 48 hours and measuring the solvent fluorescence with a fluorometer (Turner Designs Inc., San Jose, USA) (Wetzel & Likens, 2000).

#### *Temperature-standardizing dissolved oxygen concentration*

We standardized dissolved oxygen (DO) concentrations according to:

$$\mathrm{DO}_{T_{\mathrm{ref}}} = \mathrm{DO}_{\mathrm{meas}} \frac{C^*(T_{\mathrm{ref}})}{C^*(T_{\mathrm{obs}})} \tag{S3}$$

where $T_{\mathrm{ref}}$ is an arbitrary reference temperature (here, 23°C: the mean temperature in the experiment), $\mathrm{DO}_{\mathrm{meas}}$ is the dissolved oxygen concentration measured in a given tank (mg $L^{-1}$), and $C^*$ is the saturation concentration of dissolved oxygen at a given temperature (i.e., $T_{ref}$ or $T_{\mathrm{obs}}$, where pressure is 1 atmosphere and salinity is 0. $C^*$ values were calculated using the Garcia-Benson model of oxygen saturation (Garcia & Gordon, 1992) via the `o2.at.sat.base()`

function in the `LakeMetabolizer` R package (Winslow *et al.*, 2016).

#### *Estimating temperature-dependent reaeration*

To assess whether differences in air-water gas exchange rates (reaeration) at different temperatures and times of day constituted a meaningful source of bias in our dissolved oxygen estimates, we roughly estimated reaeration fluxes for each dissolved oxygen measurement based on wind speeds at each sampling time, mesocosm temperature, and gas exchange constants for fresh water.

Mean daily wind speeds for each of the six sampling dates were obtained from the Environment Canada historical climate archive (`https://climate.weather.gc.ca/historical_data`) from a nearby weather station (Vancouver International Airport, Station ID 51442). Mesocosms were significantly sheltered compared to the very exposed airport weather station site; the experiment occupied an approximately 30 x 30 m area surrounded by tall buildings and stands of coniferous trees. Mesocosms were not filled to the top, so the water surface likely experienced an additional wind buffering effect from the elevated tank edge. Mesocosms at different positions within the experimental array likely experienced similar degrees of shelter based on the distance of the nearest trees and buildings; we therefore assumed with confidence that the same wind speeds could be applied to all mesocosms on a given day. We converted wind speeds measured at the airport weather station to estimated wind speeds at the mesocosm surface on each sampling day using a shelter factor of 0.30, then used these adjusted wind speeds ($U$) to compute the gas transfer velocity normalised to turbulence index $k_{600}$ (Cole & Caraco, 1998):

$$k_{600} = 2.07 + 0.215 \times U^{1.7} \qquad \text{(S4)}$$

We then converted $k_{600}$ (eqn S4) to an $O_2$-specific gas transfer velocity constant following the Schmidt number scaling implemented in the `LakeMetabolizer` R package (Winslow *et al.*, 2016):

$$k_{O_2} = k_{600} \times \left(\frac{600}{Sc_{O_2}}\right)^{0.5} \tag{S5}$$

where the Schmidt number ($k$) for $O_2$ in freshwater was computed as a function of mesocosm temperature $T$ (°C):

$$Sc_{O_2} = 1800.6 - 120.10\,T + 3.7818\,T^2 - 0.047608\,T^3 \tag{S6}$$

Dissolved oxygen saturation concentrations were corrected to account for temperature-dependent solubility as described in the section above (Garcia & Gordon, 1992). We then estimated the oxygen saturation surplus or deficit (i.e., relative to air oxygen) for each measurement:

$$\Delta_{DO} = \mathrm{DO_{obs}} - \mathrm{DO_{sat}}(T) \tag{S7}$$

where positive values indicate supersaturation and negative values indicate undersaturation. The reaeration rate ($R_{\mathrm{reair}}$, mg $O_2$ $L^{-1}$ $day^{-1}$) for each oxygen measurement was then calculated as:

$$R_{\mathrm{reair}} = \frac{k_{O_2} \times \Delta_{DO}}{z} \tag{S8}$$

where $z$ is mesocosm depth in metres. Reaeration fluxes $F_{\mathrm{reair}}$ for each dawn-dusk-dawn measurement period were calculated by multiplying the $R_{\mathrm{reair}}$ by the time lapsed between each respective sampling. Reaeration corrections were applied to the dissolved oxygen concentrations (mg $L^{-1}$)

measured with the YSI probe prior to temperature standardisation. Corrected raw DO values at dusk and second dawn were:

$$\mathrm{DO}^{*}_{\mathrm{dusk}} = \mathrm{DO}_{\mathrm{dusk}} + F_{\mathrm{reair,dusk}} \tag{S9}$$

$$\mathrm{DO}^{*}_{\mathrm{dawn}_2} = \mathrm{DO}_{\mathrm{dawn}_2} + F_{\mathrm{reair,dawn}_2} \tag{S10}$$

We then applied a correction for temperature-dependent oxygen saturation to the reaeration-corrected dissolved oxygen values, and computed reaeration-corrected net primary production and ecosystem respiration rates, which were converted to $\mu$mol $L^{-1}$ $h^{-1}$ per metabolic theory convention. Reaeration-corrected GPP values were log-transformed and biomass-corrected, and used to estimate the model described in eqn 3. Full posterior summaries and a direct comparison of uncorrected and corrected parameter estimates are provided in Table S1.

#### *Rescaling community thermal phenotype indices*

Raw Community Thermal Phenotype Index (CTPI) as described in eqn 1 in the main text was rescaled to range from 0-1 according to:

$$\mathrm{CTPI}_{scaled} = \frac{\mathrm{CTPI}_{\mathrm{raw},c} - \min(\mathrm{CTPI}_{\mathrm{raw}})}{\max(\mathrm{CTPI}_{\mathrm{raw}}) - \min(\mathrm{CTPI}_{\mathrm{raw}})} \tag{S11}$$

Community Thermal Match Index (CTMI) as described in eqn 2 in the main text was inverted and rescaled to range from 0-1 according to:

$$\text{CTMI}_{scaled} = \frac{\max(\text{CTMI}_{\text{raw}}) - \text{CTMI}_{\text{raw},c}}{\max(\text{CTMI}_{\text{raw}}) - \min(\text{CTMI}_{\text{raw}})} \tag{S12}$$

#### *Alternative LMM test of H1 with all time points*

To test the robustness of our inferences about effects of dispersal on $E_{\text{GPP}}$ across time points, we estimated a version of the model used to test H1 (eqn 3) that included data from all time points with a three-way interaction between dispersal, temperature, and sampling week as a factor:

$$\ln(GPP \cdot M^{-1})_i = \beta_0 + E_{\text{GPP}}\,\text{Temp}_i + \beta_{T\times D_l \times W_k}\left(\text{Temp}_i \times D_{l,i} \times W_{k,i}\right) + \gamma_{metacom} + \varepsilon_i \tag{S13}$$

where $\text{Temp}_i$ is standardized Arrhenius temperature $\left(\frac{1}{kT_c} - \frac{1}{kT}\right)$ in community $i$, $M_i$ is phytoplankton community biomass, $D_{l,i}$ is a factor representing dispersal level $l \in$ {low, high} (baseline = no dispersal), $W_{k,i}$ is sampling week as a factor with $k \in \{2, 4, 6, 8, 10, 12\}$ (baseline = week 2), $\gamma_{metacom} \sim \mathcal{N}(0, \sigma^2_{\text{metacom}})$ is a metacommunity-level random intercept, and $\varepsilon_i \sim \mathcal{N}(0, \sigma^2)$ is the residual error. Posteriors were estimated using the same procedure and priors described in the Methods.

#### *Testing dispersal and temperature effects on community size spectra*

We tested whether dispersal and temperature modify phytoplankton community size spectra slopes (log abundance versus log equivalent spherical diameter) with the following model:

$$
\begin{aligned}
\ln(N_{i,c}) = \beta_0 + \beta_{\text{size}} \ln(S_{i,c}) + \beta_{\text{size}\times T} \left[\ln(S_{i,c})\,\text{Temp}_c\right] + \beta_{\text{size}\times D_l} \left[\ln(S_{i,c})\,D_{l,c}\right] \\
+ \beta_{\text{size}\times T\times D_l} \left[\ln(S_{i,c})\,\text{Temp}_c\,D_{l,c}\right] + \gamma_{metacom} + \varepsilon_{i,c}
\end{aligned}
\tag{S14}
$$

where $S_{i,c}$ is phytoplankton cell size bin (equivalent spherical diameter in 5 $\mu$m increments ranging from 0–40 $\mu$m), $N_{i,c}$ is the number of cells in size bin $i$ in community $c$, $\text{Temp}_c$ is standardized Arrhenius temperature in community $c$, $D_{l,c}$ is a factor representing dispersal level $l$ in community $c$, $\gamma_{metacom} \sim \mathcal{N}(0, \sigma^2_{\text{metacom}})$ is a metacommunity-level random intercept. Weakly informative priors were assigned (mean = 0, SD = 0.5) for all slope parameters and posteriors were estimated with the same procedure described in the Methods. Full results are reported in Table S7 and Figure S12.

#### *Testing effect of dispersal on Oocystis relative abundance*

To test whether dispersal changed the relative abundance of *Oocystis* sp., we modelled its logit-transformed relative abundance as:

$$
\text{logit}(p_i) = \beta_0 + \beta_{D_l} D_{l,i} + \beta_{W_k} W_{k,i} + \beta_{D_l\times W_k} (D_{l,i} \times W_{k,i}) + \beta_T T_i + \gamma_{metacom} + \varepsilon_i \tag{S15}
$$

where $p_i$ is the relative abundance of *Oocystis* sp. in mesocosm (community) $i$, $D_{l,i}$ is a factor representing dispersal level $l \in$ {low, high} (baseline = no dispersal); $W_{k,i}$ is sampling week as a factor with $k \in \{4, 8, 12\}$ (baseline = week 4), $D_{l,i} \times W_{k,i}$ are their interactions, $T_i$ is temperature in mesocosm $i$, $\gamma_m \sim \mathcal{N}(0, \sigma^2_{\text{meta}})$ is a metacommunity-level random intercept, and $\varepsilon_i \sim \mathcal{N}(0, \sigma^2)$ is the residual error. Weakly informative priors were assigned (mean = 0, SD = 0.5) for all slope parameters and posteriors were estimated with the same procedure described in the Methods. Results are shown in Figure S15.

### Supplementary Tables

**Table S1:** Comparison of $E_{GPP}$ posterior estimates from versions of the LMM test of Hypothesis 1 to uncorrected (as in manuscript) and reaeration-corrected GPP estimates. Bold values indicate posterior coefficient estimates where < 5% of the 89% HDI falls within the Region of Practical Equivalence (ROPE, defined as $\pm 0.20 \times$ SD(GPP)), i.e. the effect is judged nonzero. Bias = corrected − uncorrected posterior mean.

| **Week** | **Parameter** | **Uncorrected mean [89% HDI]** | **Corrected mean [89% HDI]** | $\boldsymbol{E_{GPP}}$ **bias (eV)** |
|---|---|---|---|---|
| Week 2 | $E_{GPP}$ (none) | 0.16 [-0.19, 0.51] | 0.29 [-0.10, 0.68] | +0.129 |
| | $E_{GPP}$ (low) | 0.09 [-0.47, 0.65] | 0.06 [-0.63, 0.77] | -0.023 |
| | $E_{GPP}$ (high) | -0.05 [-0.61, 0.52] | -0.02 [-0.70, 0.68] | +0.031 |
| | $\Delta E_{GPP}$ (low − none) | -0.07 [-0.63, 0.49] | -0.23 [-0.88, 0.43] | -0.152 |
| | $\Delta E_{GPP}$ (high − none) | -0.21 [-0.76, 0.34] | -0.30 [-0.94, 0.34] | -0.097 |
| Week 4 | $E_{GPP}$ (none) | -0.04 [-0.29, 0.23] | 0.28 [-0.07, 0.65] | +0.321 |
| | $E_{GPP}$ (low) | 0.02 [-0.34, 0.40] | -0.07 [-0.69, 0.56] | -0.088 |
| | $E_{GPP}$ (high) | 0.11 [-0.23, 0.44] | 0.14 [-0.45, 0.74] | +0.034 |
| | $\Delta E_{GPP}$ (low − none) | 0.06 [-0.36, 0.46] | -0.35 [-0.95, 0.24] | -0.408 |
| | $\Delta E_{GPP}$ (high − none) | 0.14 [-0.24, 0.53] | -0.14 [-0.73, 0.45] | -0.287 |
| Week 6 | $E_{GPP}$ (none) | 0.29 [0.01, 0.57] | **0.29 [0.04, 0.55]** | +0.004 |
| | $E_{GPP}$ (low) | **0.40 [0.10, 0.69]** | **0.35 [0.07, 0.64]** | -0.048 |
| | $E_{GPP}$ (high) | 0.24 [-0.07, 0.54] | 0.25 [-0.05, 0.54] | +0.011 |
| | $\Delta E_{GPP}$ (low − none) | 0.11 [-0.17, 0.38] | 0.06 [-0.21, 0.32] | -0.052 |
| | $\Delta E_{GPP}$ (high − none) | -0.05 [-0.35, 0.23] | -0.04 [-0.33, 0.22] | +0.006 |
| Week 8 | $E_{GPP}$ (none) | **0.65 [0.43, 0.88]** | **0.56 [0.34, 0.78]** | -0.093 |
| | $E_{GPP}$ (low) | **0.56 [0.30, 0.81]** | **0.53 [0.27, 0.78]** | -0.032 |
| | $E_{GPP}$ (high) | 0.17 [-0.09, 0.44] | 0.16 [-0.09, 0.42] | -0.012 |
| | $\Delta E_{GPP}$ (low − none) | -0.09 [-0.36, 0.16] | -0.03 [-0.30, 0.22] | +0.061 |
| | $\Delta E_{GPP}$ (high − none) | **-0.48 [-0.75, -0.21]** | **-0.40 [-0.67, -0.14]** | +0.082 |
| Week 10 | $E_{GPP}$ (none) | **0.72 [0.44, 0.99]** | **0.69 [0.40, 0.97]** | -0.033 |

*Table S1 (continued)*

| Week | Parameter | Uncorrected mean [89% HDI] | Corrected mean [89% HDI] | $E_{GPP}$ bias (eV) |
|---|---|---|---|---|
| | $E_{GPP}$ (low) | **0.53 [0.13, 0.92]** | **0.60 [0.16, 1.03]** | +0.075 |
| | $E_{GPP}$ (high) | **0.79 [0.39, 1.19]** | **0.70 [0.27, 1.12]** | -0.098 |
| | $\Delta E_{GPP}$ (low − none) | -0.19 [-0.62, 0.23] | -0.09 [-0.54, 0.37] | +0.109 |
| | $\Delta E_{GPP}$ (high − none) | 0.07 [-0.38, 0.52] | 0.01 [-0.46, 0.47] | -0.065 |
| Week 12 | $E_{GPP}$ (none) | **0.42 [0.14, 0.71]** | **0.38 [0.09, 0.67]** | -0.038 |
| | $E_{GPP}$ (low) | **0.49 [0.12, 0.86]** | **0.39 [0.03, 0.76]** | -0.099 |
| | $E_{GPP}$ (high) | **0.67 [0.30, 1.03]** | **0.55 [0.19, 0.92]** | -0.113 |
| | $\Delta E_{GPP}$ (low − none) | 0.07 [-0.25, 0.39] | 0.01 [-0.30, 0.33] | -0.062 |
| | $\Delta E_{GPP}$ (high − none) | 0.24 [-0.08, 0.55] | 0.17 [-0.15, 0.47] | -0.075 |

**Table S2:** List of LMMs for each response variable in the SEM of direct and indirect effects on Gross Primary Productivity. All path coefficients are allowed to vary by sampling week via interactions with week as a factor ($W_k$, baseline = week 4). Phytoplankton community size spectra, phytoplankton community thermal phenotype composition (community thermal phenotype index), phytoplankton biomass, and zooplankton biomass mediate indirect effects of dispersal on GPP. Temperature, dispersal, and total N are exogenous variables.

| Response | Model |
|---|---|
| Size spectra ($S$) | $S_i = \beta_{0,S} + (\beta_{S,T}\, T_i + \beta_{S,Z}\, Z_i + \beta_{S,D_{\text{low}}}\, D_{\text{low}} + \beta_{S,D_{\text{high}}}\, D_{\text{high}}) \times W_k + u_S + \varepsilon_S$ |
| Thermal traits ($C$) | $C_i = \beta_{0,C} + (\beta_{C,T}\, T_i + \beta_{C,Z}\, Z_i + \beta_{C,D_{\text{low}}}\, D_{\text{low}} + \beta_{C,D_{\text{high}}}\, D_{\text{high}}) \times W_k + u_C + \varepsilon_C$ |
| Phyto biomass ($B$) | $B_i = \beta_{0,B} + (\beta_{B,T}\, T_i + \beta_{B,N}\, N_i + \beta_{B,Z}\, Z_i + \beta_{B,D_{\text{low}}}\, D_{\text{low}} + \beta_{B,D_{\text{high}}}\, D_{\text{high}}) \times W_k + u_B + \varepsilon_B$ |
| Zoop biomass ($Z$) | $Z_i = \beta_{0,Z} + (\beta_{Z,T}\, T_i + \beta_{Z,D_{\text{low}}}\, D_{\text{low}} + \beta_{Z,D_{\text{high}}}\, D_{\text{high}}) \times W_k + u_Z + \varepsilon_Z$ |
| GPP | $GPP_i = \beta_{0,G} + (\beta_{G,S}\, S_i + \beta_{G,C}\, C_i + \beta_{G,B}\, B_i + \beta_{G,T}\, T_i + \beta_{G,N}\, N_i) \times W_k + u_{GPP} + \varepsilon_{GPP}$ |

$S_i$ = size spectra slope, $C_i$ = community thermal phenotype index, $B_i$ = ln phytoplankton biomass, $Z_i$ = ln zooplankton biomass, $T_i$ = temperature ($\frac{1}{kT_c} - \frac{1}{kT}$), $N_i$ = ln total nitrogen ($\mu$M), $D_{\text{low}}$ and $D_{\text{high}}$ are categorical dispersal levels (baseline = no dispersal), $W_k$ = sampling week as a factor with $k \in \{4, 8, 12\}$ (baseline = week 4). $u$ = random intercept for metacommunity, $u \sim \mathcal{N}(0, \sigma_{\text{meta}})$. Residuals $\varepsilon \sim \mathcal{N}(0, \sigma)$.

**Table S3:** Full summary of SEM path posteriors (mean coefficient estimate, 89% highest density interval (HDI), and proportion of the 89% HDI within ROPE = ±0.20×SD(GPP), computed using the pooled standard deviation across all sampling weeks). Week enters the model as a moderator of all path coefficients; week-specific estimates are recovered as the sum of the main effect and the corresponding week interaction term (baseline = week 4). Bold week-specific rows indicate HDI within ROPE < 5% (non-zero effect), and bold path names indicate those with non-zero effects across all three weeks.

| Path | Week | Coefficient | 89% HDI | % HDI in ROPE |
|---|---|---|---|---|
| Temp → GPP | **4** | **0.309** | **[0.152, 0.470]** | **0.00** |
| | **8** | **0.579** | **[0.413, 0.743]** | **0.00** |
| | **12** | **0.665** | **[0.514, 0.813]** | **0.00** |
| Total N → GPP | 4 | 0.015 | [-0.011, 0.041] | 100.00 |
| | 8 | −0.002 | [-0.028, 0.024] | 100.00 |
| | 12 | 0.006 | [-0.024, 0.035] | 100.00 |
| Size spectra → GPP | 4 | −0.037 | [-0.085, 0.011] | 55.30 |
| | 8 | −0.046 | [-0.099, 0.007] | 42.80 |
| | 12 | −0.089 | [-0.172, −0.005] | 21.50 |
| Thermal phenotypes → GPP | 4 | 0.014 | [-0.100, 0.131] | 36.50 |
| | 8 | −0.009 | [-0.151, 0.130] | 30.00 |
| | 12 | 0.065 | [-0.193, 0.322] | 16.30 |
| Phyto biomass → GPP | 4 | 0.142 | [0.007, 0.276] | 11.60 |
| | **8** | **0.366** | **[0.249, 0.480]** | **0.00** |
| | 12 | 0.117 | [0.020, 0.216] | 13.10 |
| **Temp → Zoop biomass** | **4** | **0.987** | **[0.413, 1.550]** | **0.00** |
| | **8** | **0.918** | **[0.159, 1.680]** | **0.00** |
| | **12** | **1.270** | **[0.548, 1.960]** | **0.00** |
| Temp → Size spectra | 4 | −0.238 | [-0.690, 0.203] | 27.40 |
| | 8 | −0.471 | [-1.040, 0.102] | 17.50 |
| | 12 | 0.197 | [-0.332, 0.730] | 12.20 |
| Temp → Thermal phenotypes | 4 | 0.271 | [-0.002, 0.541] | 6.42 |
| | **8** | **0.336** | **[0.057, 0.617]** | **0.00** |

*Table S3 (continued)*

| Path | Week | Coefficient | 89% HDI | % HDI in ROPE |
|---|---|---|---|---|
| | 12 | 0.233 | [-0.026, 0.490] | 6.75 |
| Temp → Phyto biomass | **4** | **0.353** | **[0.057, 0.643]** | **0.00** |
| | 8 | 0.181 | [-0.124, 0.488] | 15.80 |
| | **12** | **0.296** | **[0.016, 0.571]** | **4.70** |
| Total N → Phyto biomass | 4 | −0.010 | [-0.063, 0.042] | 79.30 |
| | 8 | −0.002 | [-0.057, 0.053] | 76.40 |
| | 12 | 0.008 | [-0.050, 0.068] | 71.00 |
| Disp: low → Zoop biomass | 4 | −0.113 | [-0.540, 0.313] | 45.70 |
| | 8 | −0.042 | [-0.547, 0.479] | 23.90 |
| | 12 | −0.073 | [-0.580, 0.434] | 43.20 |
| Disp: high → Zoop biomass | 4 | 0.075 | [-0.347, 0.496] | 46.40 |
| | 8 | 0.260 | [-0.235, 0.771] | 24.40 |
| | 12 | 0.169 | [-0.347, 0.696] | 42.00 |
| Disp: low → Size spectra | 4 | 0.203 | [-0.073, 0.475] | 36.20 |
| | 8 | −0.241 | [-0.533, 0.049] | 25.20 |
| | 12 | 0.059 | [-0.243, 0.359] | 21.50 |
| Disp: high → Size spectra | 4 | −0.277 | [-0.543, −0.009] | 19.40 |
| | 8 | −0.189 | [-0.487, 0.108] | 33.50 |
| | 12 | −0.046 | [-0.356, 0.262] | 20.90 |
| Disp: low → Thermal phenotypes | 4 | 0.029 | [-0.104, 0.161] | 34.50 |
| | 8 | 0.083 | [-0.039, 0.211] | 32.70 |
| | 12 | 0.093 | [-0.037, 0.226] | 17.20 |
| Disp: high → Thermal phenotypes | 4 | 0.126 | [0.000, 0.256] | 18.60 |
| | 8 | 0.124 | [-0.002, 0.252] | 16.40 |
| | 12 | 0.050 | [-0.084, 0.183] | 17.00 |
| Disp: low → Phyto biomass | 4 | 0.153 | [-0.013, 0.313] | 17.30 |
| | 8 | 0.144 | [-0.020, 0.304] | 17.30 |

*Table S3 (continued)*

| Path | Week | Coefficient | 89% HDI | % HDI in ROPE |
|---|---|---|---|---|
| | 12 | 0.072 | [-0.095, 0.237] | 25.30 |
| Disp: high → Phyto biomass | 4 | 0.072 | [-0.085, 0.231] | 26.60 |
| | **8** | **0.311** | **[0.149, 0.474]** | **0.00** |
| | **12** | **0.223** | **[0.060, 0.385]** | **0.00** |
| Zoop biomass → Size spectra | 4 | −0.114 | [-0.245, 0.015] | 51.60 |
| | 8 | 0.042 | [-0.156, 0.239] | 50.40 |
| | 12 | −0.025 | [-0.145, 0.092] | 54.40 |
| Zoop biomass → Therm phenotypes | 4 | −0.010 | [-0.063, 0.045] | 83.70 |
| | 8 | −0.021 | [-0.101, 0.059] | 52.20 |
| | 12 | 0.004 | [-0.044, 0.051] | 47.40 |
| Zoop biomass → Phyto biomass | 4 | −0.043 | [-0.101, 0.016] | 43.90 |
| | 8 | 0.001 | [-0.087, 0.091] | 54.50 |
| | 12 | 0.035 | [-0.018, 0.087] | 69.10 |

**Table S4:** Structural Equation Model (SEM) posterior sensitivity to ROPE width. The proportion of the posterior 89% HDI is reported for three ROPE widths (5%, 10%, or 20% of the pooled standard deviation of GPP across all weeks) for each SEM path and week. Week-specific estimates were estimated from the sum of the main effect and the corresponding week interaction term (baseline = week 4). Bold entries indicate the specific week×path combinations that are sensitive to ROPE width.

| Path | Week | ROPE = ±0.05×SD | ROPE = ±0.10×SD | ROPE = ±0.20×SD |
|---|---|---|---|---|
| Temp → GPP (direct) | 4 | 0.00 | 0.00 | 0.00 |
| | 8 | 0.00 | 0.00 | 0.00 |
| | 12 | 0.00 | 0.00 | 0.00 |
| Temp → GPP (indirect) | 4 | 13.6 | 20.6 | 34.6 |
| | 8 | 8.00 | 16.0 | 31.9 |
| | 12 | 10.3 | 20.7 | 41.3 |
| High dispersal → GPP (indirect) | 4 | 29.4 | 48.3 | 77.7 |
| | 8 | 0.00 | 0.00 | 0.00 |
| | 12 | 20.8 | 37.8 | 58.6 |
| Low dispersal → GPP (indirect) | 4 | 26.9 | 53.7 | 86.5 |
| | 8 | 10.3 | 18.3 | 34.2 |
| | 12 | 20.9 | 41.7 | 83.4 |
| Total N → GPP | 4 | 41.1 | 62.5 | 100.0 |
| | 8 | 40.4 | 80.8 | 100.0 |
| | 12 | 35.8 | 71.6 | 100.0 |
| **Size spectra → GPP** | 4 | 21.8 | 33.5 | 55.3 |
| | 8 | 13.1 | 23.0 | 42.8 |
| | **12** | **2.50** | **8.82** | **21.5** |
| **Thermal traits → GPP** | 4 | 9.12 | 18.2 | 36.5 |
| | 8 | 7.49 | 15.0 | 30.0 |
| | **12** | **4.08** | **8.17** | **16.3** |
| **Phyto biomass → GPP** | **4** | **0.00** | **3.73** | **11.6** |
| | 8 | 0.00 | 0.00 | 0.00 |

*Table S4 (continued)*

| Path | Week | ROPE = ±0.05× SD | ROPE = ±0.10× SD | ROPE = ±0.20× SD |
|---|---|---|---|---|
| | **12** | **0.00** | **2.39** | **13.1** |
| Temp → Zoop biomass | 4 | 0.00 | 0.00 | 0.00 |
| | 8 | 0.00 | 0.00 | 0.00 |
| | 12 | 0.00 | 0.00 | 0.00 |
| **Temp → Size spectra** | **4** | **2.36** | **4.72** | **9.44** |
| | **8** | **1.85** | **3.71** | **7.41** |
| | **12** | **1.98** | **3.97** | **7.94** |
| **Temp → Thermal traits** | **4** | **0.00** | **1.89** | **5.78** |
| | 8 | 0.00 | 0.00 | 0.00 |
| | **12** | **4.10** | **6.44** | **10.5** |
| **Temp → Phyto biomass** | 4 | 0.00 | 0.00 | 0.00 |
| | **8** | **3.44** | **6.89** | **13.8** |
| | 12 | 0.00 | 0.00 | 2.67 |
| Total N → Phyto biomass | 4 | 20.1 | 40.2 | 79.3 |
| | 8 | 19.1 | 38.2 | 76.4 |
| | 12 | 17.7 | 35.5 | 71.0 |
| **Disp: low → Zoop biomass** | **4** | **2.47** | **4.94** | **9.88** |
| | **8** | **2.05** | **4.10** | **8.21** |
| | **12** | **2.08** | **4.15** | **8.31** |
| **Disp: high → Zoop biomass** | **4** | **2.51** | **5.02** | **10.0** |
| | **8** | **2.10** | **4.19** | **8.38** |
| | **12** | **2.02** | **4.04** | **8.08** |
| **Disp: low → Size spectra** | **4** | **3.85** | **7.69** | **15.4** |
| | **8** | **3.62** | **7.24** | **14.5** |
| | **12** | **3.50** | **7.00** | **14.0** |
| **Disp: high → Size spectra** | 4 | 0.00 | 0.38 | 4.33 |

*Table S4 (continued)*

| **Path** | **Week** | **ROPE = ±0.05× SD** | **ROPE = ±0.10× SD** | **ROPE = ±0.20× SD** |
|---|---|---|---|---|
| | **8** | **3.54** | **7.09** | **14.2** |
| | **12** | **3.42** | **6.83** | **13.7** |
| Disp: low → Thermal traits | 4 | 7.97 | 15.9 | 31.9 |
| | 8 | 8.42 | 16.8 | 33.0 |
| | 12 | 8.01 | 16.0 | 30.5 |
| **Disp: high → Thermal traits** | **4** | **4.85** | **8.97** | **17.2** |
| | **8** | **4.17** | **8.33** | **16.6** |
| | 12 | 7.91 | 15.8 | 31.6 |
| Disp: low → Phyto biomass | 4 | 6.47 | 10.8 | 17.3 |
| | 8 | 6.50 | 10.8 | 17.3 |
| | 12 | 6.32 | 12.6 | 25.3 |
| Disp: high → Phyto biomass | 4 | 6.66 | 13.3 | 26.6 |
| | 8 | 0.00 | 0.00 | 0.00 |
| | 12 | 0.00 | 0.00 | 0.00 |
| Zoop biomass → Size spectra | 4 | 8.17 | 12.3 | 20.4 |
| | 8 | 5.33 | 10.7 | 21.3 |
| | 12 | 8.87 | 17.7 | 35.5 |
| Zoop biomass → Thermal traits | 4 | 19.7 | 39.4 | 78.9 |
| | 8 | 13.2 | 26.5 | 52.9 |
| | 12 | 22.0 | 44.1 | 88.1 |
| Zoop biomass → Phyto biomass | 4 | 18.0 | 30.3 | 48.3 |
| | 8 | 11.9 | 23.8 | 47.6 |
| | 12 | 20.0 | 38.4 | 58.4 |

**Table S5:** ANOVA and Tukey HSD post-hoc tests for within-metacommunity coefficients of variation (CV) of biomass, size spectra, and community thermal phenotype index (CTPI) across weeks and dispersal treatments. Bold rows indicate Tukey HSD $p < 0.05$.

| **Trait** | **Week** | $F_{2,9}$ | $p_{\textbf{ANOVA}}$ | **Contrast** | **Estimate** | $p_{\textbf{Tukey}}$ |
|---|---|---|---|---|---|---|
| Biomass CV | 2 | 0.17 | 0.85 | none - low | 0.00 | 0.99 |
| | | | | none - high | 0.02 | 0.84 |
| | | | | low - high | 0.01 | 0.91 |
| | 4 | 1.91 | 0.20 | none - low | 0.05 | 0.23 |
| | | | | none - high | 0.04 | 0.30 |
| | | | | low - high | 0.00 | 0.98 |
| | 6 | 1.82 | 0.22 | none - low | -0.03 | 0.23 |
| | | | | none - high | -0.01 | 0.95 |
| | | | | low - high | 0.03 | 0.34 |
| | 8 | 2.90 | 0.11 | none - low | 0.01 | 0.67 |
| | | | | none - high | 0.03 | 0.09 |
| | | | | low - high | 0.02 | 0.33 |
| | 10 | 1.87 | 0.21 | none - low | 0.03 | 0.71 |
| | | | | none - high | 0.06 | 0.19 |
| | | | | low - high | 0.04 | 0.53 |
| | 12 | 3.06 | 0.10 | none - low | 0.01 | 0.97 |
| | | | | none - high | 0.05 | 0.12 |
| | | | | low - high | 0.05 | 0.16 |
| Size spectra CV | **2** | **10.04** | **0.01** | **none - low** | **0.19** | **0.01** |
| | | | | **none - high** | **0.21** | **0.01** |
| | | | | low - high | 0.02 | 0.88 |
| | 4 | 1.91 | 0.20 | none - low | 0.05 | 0.94 |
| | | | | none - high | 0.25 | 0.21 |
| | | | | low - high | 0.21 | 0.33 |
| | 6 | 0.40 | 0.68 | none - low | 0.06 | 0.77 |

*Table S5 (continued)*

| **Trait** | **Week** | $F_{2,9}$ | $p_{\textbf{ANOVA}}$ | **Contrast** | **Estimate** | $p_{\textbf{Tukey}}$ |
|---|---|---|---|---|---|---|
| | | | | none - high | 0.07 | 0.70 |
| | | | | low - high | 0.01 | 0.99 |
| | 8 | 0.27 | 0.77 | none - low | 0.03 | 0.91 |
| | | | | none - high | -0.03 | 0.94 |
| | | | | low - high | -0.06 | 0.75 |
| | 10 | 1.48 | 0.28 | none - low | 0.00 | 1.00 |
| | | | | none - high | 0.08 | 0.34 |
| | | | | low - high | 0.08 | 0.34 |
| | 12 | 1.32 | 0.32 | none - low | -0.01 | 0.91 |
| | | | | none - high | 0.04 | 0.51 |
| | | | | low - high | 0.05 | 0.31 |
| CTMI CV | 4 | 0.35 | 0.71 | none - low | 0.04 | 0.99 |
| | | | | none - high | 0.23 | 0.72 |
| | | | | low - high | 0.19 | 0.80 |
| | 8 | 0.21 | 0.81 | none - low | 0.07 | 0.93 |
| | | | | none - high | -0.05 | 0.96 |
| | | | | low - high | -0.11 | 0.80 |
| | 12 | 0.01 | 0.99 | none - low | -0.01 | 1.00 |
| | | | | none - high | 0.00 | 1.00 |
| | | | | low - high | 0.01 | 1.00 |

**Table S6:** Levene's tests for equality of variances in GPP and phytoplankton community biomass among dispersal treatments. $p < 0.05$ are shown in bold.

| | **GPP** | | | **Biomass** | | |
|---|---|---|---|---|---|---|
| Week | df | *F* | *p* | df | *F* | *p* |
| 2 | 2, 44 | 2.99 | 0.06 | 2, 45 | 0.53 | 0.59 |
| 4 | 2, 42 | 2.02 | 0.15 | 2, 45 | 2.09 | 0.14 |
| 6 | 2, 45 | 0.23 | 0.79 | 2, 45 | 0.55 | 0.58 |
| 8 | 2, 43 | 8.12 | **0.0010** | 2, 45 | 3.12 | 0.054 |
| 10 | 2, 45 | 4.25 | **0.020** | 2, 45 | 2.50 | 0.093 |
| 12 | 2, 43 | 0.94 | 0.40 | 2, 45 | 4.46 | **0.017** |

**Table S7:** Linear mixed effects model describing dispersal and temperature effects on phytoplankton community size spectra (log abundance versus log equivalent spherical diameter). The model structure is described in eqn S14.

| Week | Term | Mean | HDI Low | HDI High | % HDI in ROPE |
|---|---|---|---|---|---|
| **2** | **Size slope** | **-5.24** | **-7.17** | **-3.27** | **0** |
| | High dispersal × Size | 1.39 | -1.46 | 3.80 | 6.73 |
| | **Low dispersal × Size** | **3.02** | **0.39** | **5.80** | **0** |
| | Size × Temp | 0.13 | 0.05 | 0.22 | 75.7 |
| | High dispersal × Size × Temp | -0.07 | -0.18 | 0.05 | 100 |
| | Low dispersal × Size × Temp | -0.13 | -0.24 | -0.01 | 72.1 |
| **4** | **Size slope** | **-3.10** | **-5.03** | **-1.10** | **0** |
| | High dispersal × Size | 0.47 | -2.09 | 3.10 | 6.82 |
| | Low dispersal × Size | 1.22 | -1.64 | 4.11 | 6.16 |
| | Size × Temp | 0.06 | -0.02 | 0.14 | 100 |
| | High dispersal × Size × Temp | -0.02 | -0.13 | 0.09 | 100 |
| | Low dispersal × Size × Temp | -0.04 | -0.16 | 0.08 | 100 |
| **6** | **Size slope** | **-2.33** | **-3.73** | **-0.97** | **0** |
| | **High dispersal × Size** | **-1.99** | **-3.81** | **-0.08** | **2.6** |
| | Low dispersal × Size | -0.92 | -2.89 | 1.03 | 9.04 |
| | Size × Temp | 0.01 | -0.04 | 0.06 | 100 |
| | High dispersal × Size × Temp | 0.07 | 0.00 | 0.14 | 100 |
| | Low dispersal × Size × Temp | 0.03 | -0.04 | 0.10 | 100 |
| **8** | Size slope | 0.44 | -0.92 | 1.80 | 13.0 |
| | **High dispersal × Size** | **-3.24** | **-5.33** | **-1.18** | **0** |
| | Low dispersal × Size | -0.72 | -3.07 | 1.43 | 7.87 |
| | Size × Temp | -0.08 | -0.13 | -0.03 | 100 |
| | High dispersal × Size × Temp | 0.12 | 0.05 | 0.20 | 85.4 |
| | Low dispersal × Size × Temp | 0.02 | -0.06 | 0.11 | 100 |
| **10** | Size slope | 0.90 | -0.32 | 2.22 | 14.0 |
| | **High dispersal × Size** | **-3.53** | **-5.31** | **-1.76** | **0** |
| | **Low dispersal × Size** | **-2.59** | **-4.40** | **-0.75** | **0** |
| | Size × Temp | -0.13 | -0.19 | -0.08 | 89.5 |
| | High dispersal × Size × Temp | 0.15 | 0.07 | 0.22 | 69.3 |
| | Low dispersal × Size × Temp | 0.11 | 0.03 | 0.19 | 92.4 |
| **12** | **Size slope** | **-3.78** | **-4.99** | **-2.43** | **0** |
| | High dispersal × Size | -0.23 | -2.00 | 1.52 | 10.1 |
| | **Low dispersal × Size** | **2.02** | **0.26** | **3.72** | **0** |
| | Size × Temp | 0.06 | 0.00 | 0.12 | 100 |
| | High dispersal × Size × Temp | 0.02 | -0.06 | 0.11 | 100 |
| | Low dispersal × Size × Temp | -0.09 | -0.17 | -0.00 | 100 |

### Supplementary Figures

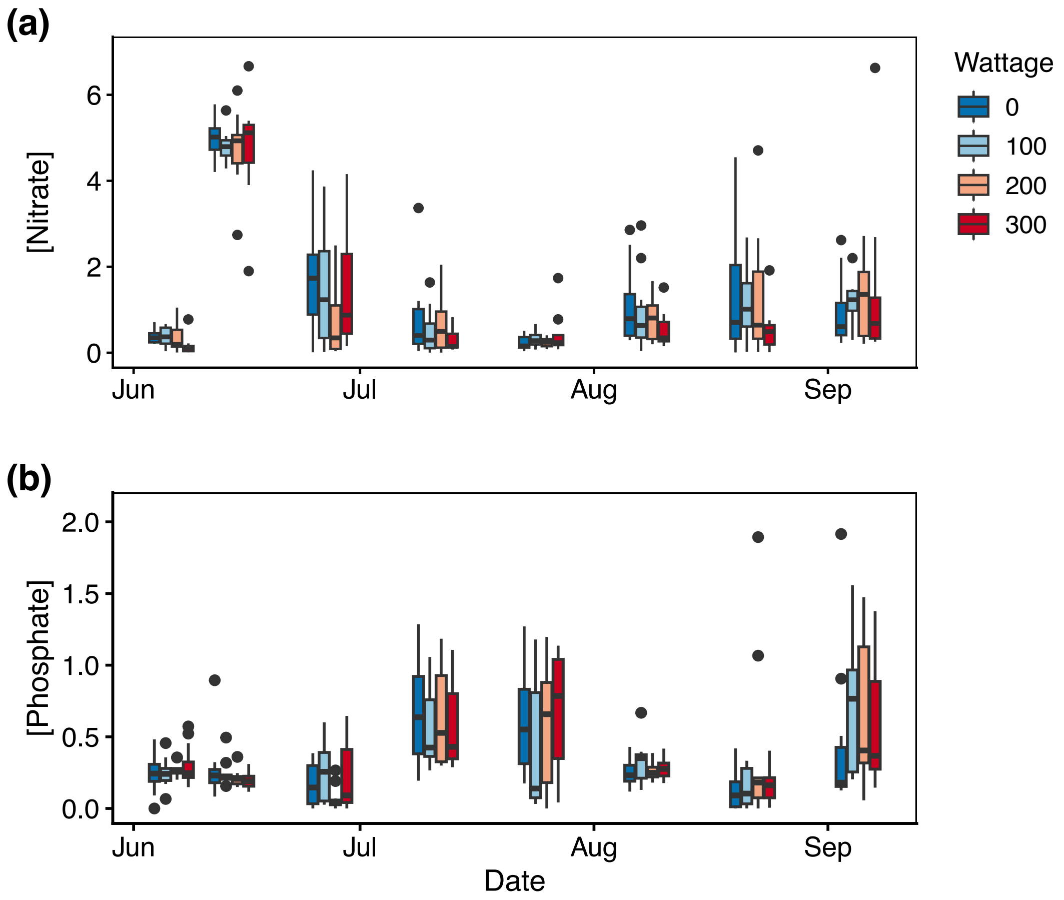


**Figure S1:** Water column **(a)** nitrate and **(b)** phosphate concentrations ($\mu$M) over the 12-week experiment. Colours indicate warming treatment.

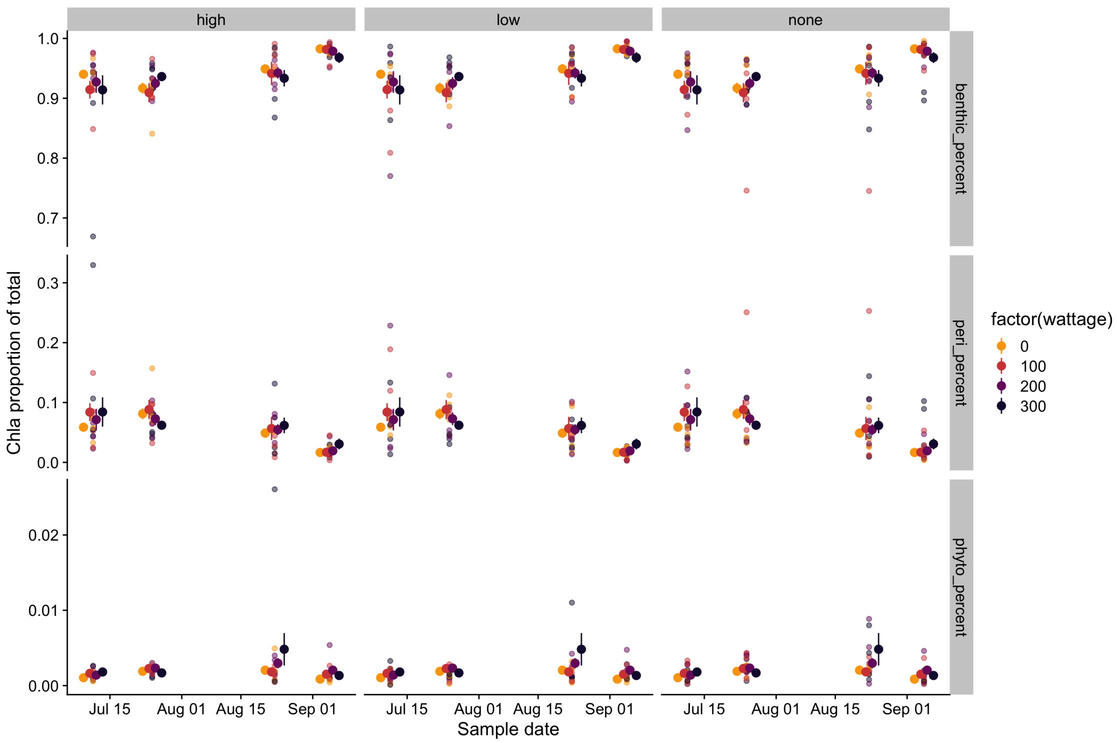


**Figure S2:** Chlorophyll concentrations of different autotroph communities (phytoplankton, periphyton, benthic microalgae) within all mesocosms over time. Proportions of these groups did not differ significantly among experimental treatments.

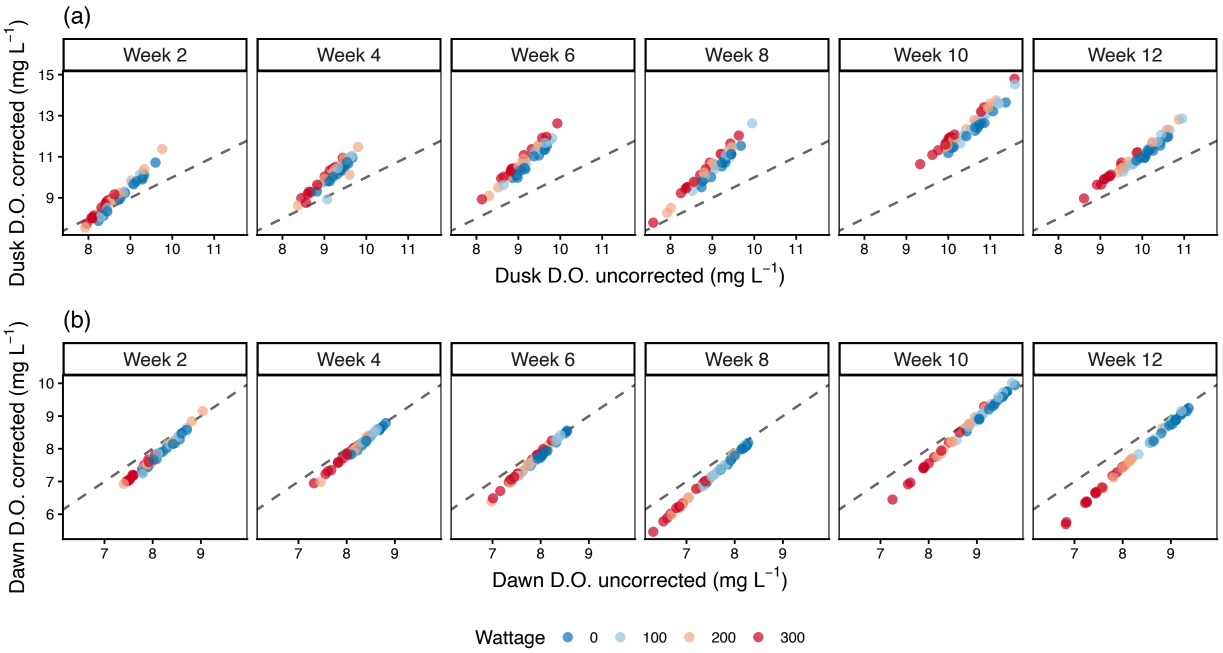


**Figure S3:** Visualizing re-aeration (air-water $O_2$ exchange) correction effects on (a) dusk and (b) dawn dissolved oxygen measurements. Dashed line represents the 1:1 line, where points above the line suggest uncorrected values may be under-estimates of true dissolved oxygen concentration, and points below the line suggest uncorrected values may be over-estimates. The calculation used to perform the reaeration correction is described in *Estimating temperature-dependent reaeration*.

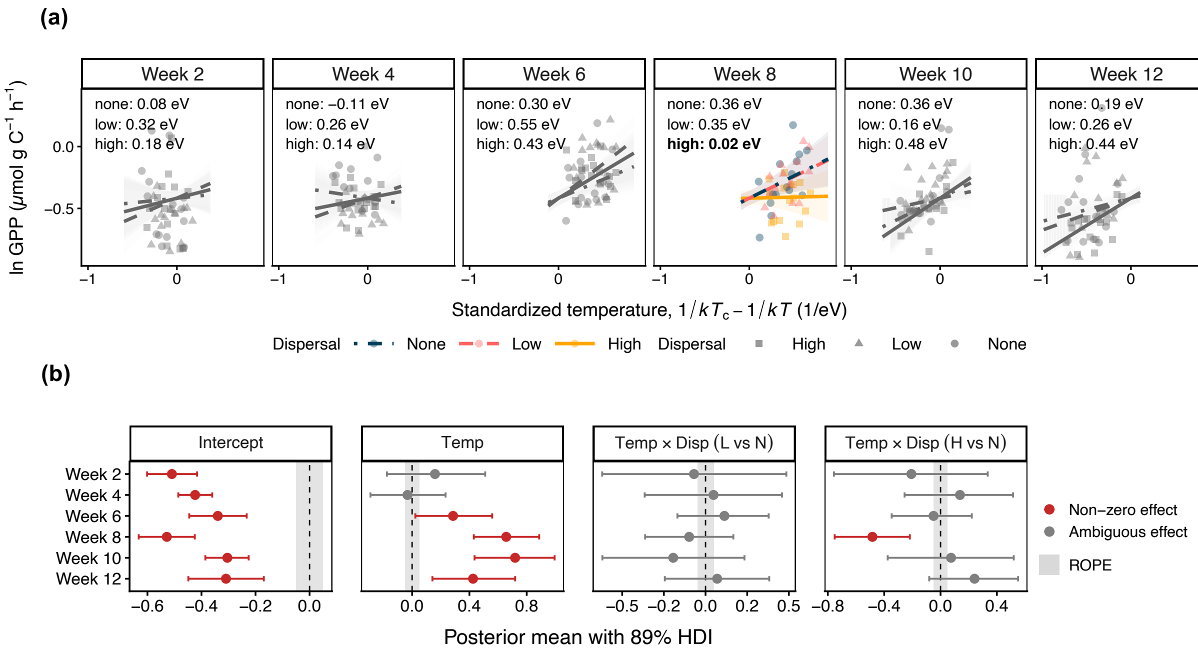


**Figure S4:** Results for an alternative version of the LMM test of H1 using data from all sampling weeks and with a three-way temperature × dispersal × week interaction. **(a)** Coloured points and lines indicate weeks where dispersal modified the slope ($E_{\mathrm{GPP}}$) of the temperature-GPP relationship. Grey lines indicate weeks where $E_{\mathrm{GPP}}$ did not significantly differ among dispersal treatments. **(b)** Forest plot of posterior estimates (± 89% HDI) for each fixed effect in the GLMMs represented in panel (a). Grey shaded regions represent Region of Practical Equivalence. The "Temp × Disp" panels represent pairwise differences in the posterior slope for temperature versus GPP ($E_{\mathrm{GPP}}$) among the three dispersal treatments for a given week, inferred from the "temp:disp:week" interaction term.

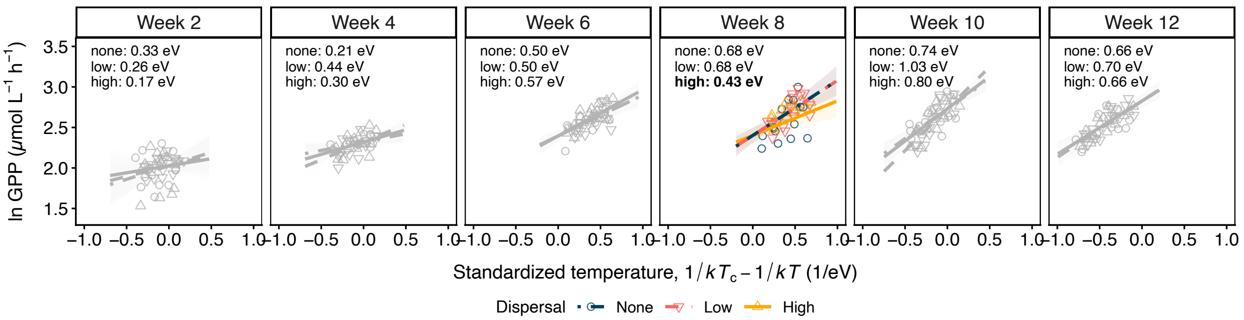


**Figure S5:** Alternative version of model represented in figure 3 using raw GPP ($\mu$mol L$^{-1}$ h$^{-1}$) instead of biomass-corrected GPP. Coloured points and lines indicate weeks where dispersal modified the slope ($E_{\mathrm{GPP}}$) of the temperature-GPP relationship. Grey lines indicate weeks where $E_{\mathrm{GPP}}$ did not significantly differ among dispersal treatments.

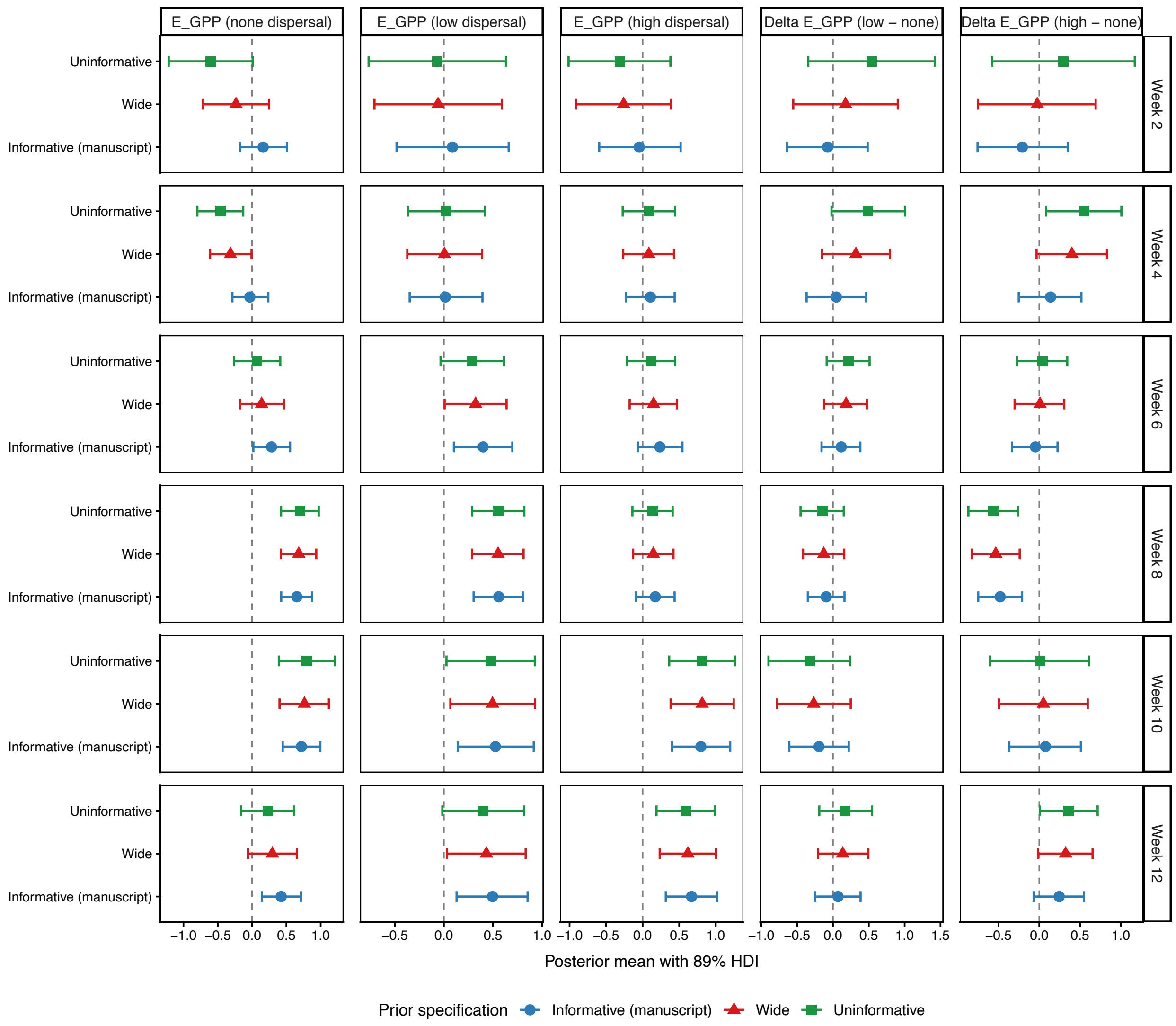


**Figure S6:** Prior sensitivity analysis for LMM test of H1. Panels compare posterior estimates (± 89% HDI) from estimating the model described in equation 3 using informative priors as in the manuscript (mean = 0.65, SD = 0.28; blue points), wide informative priors with a twice as large standard deviation (mean = 0.65, SD = 0.6; red points), and uninformative priors (mean = 0, SD = 10; green points) for $E_{\text{GPP}}$.

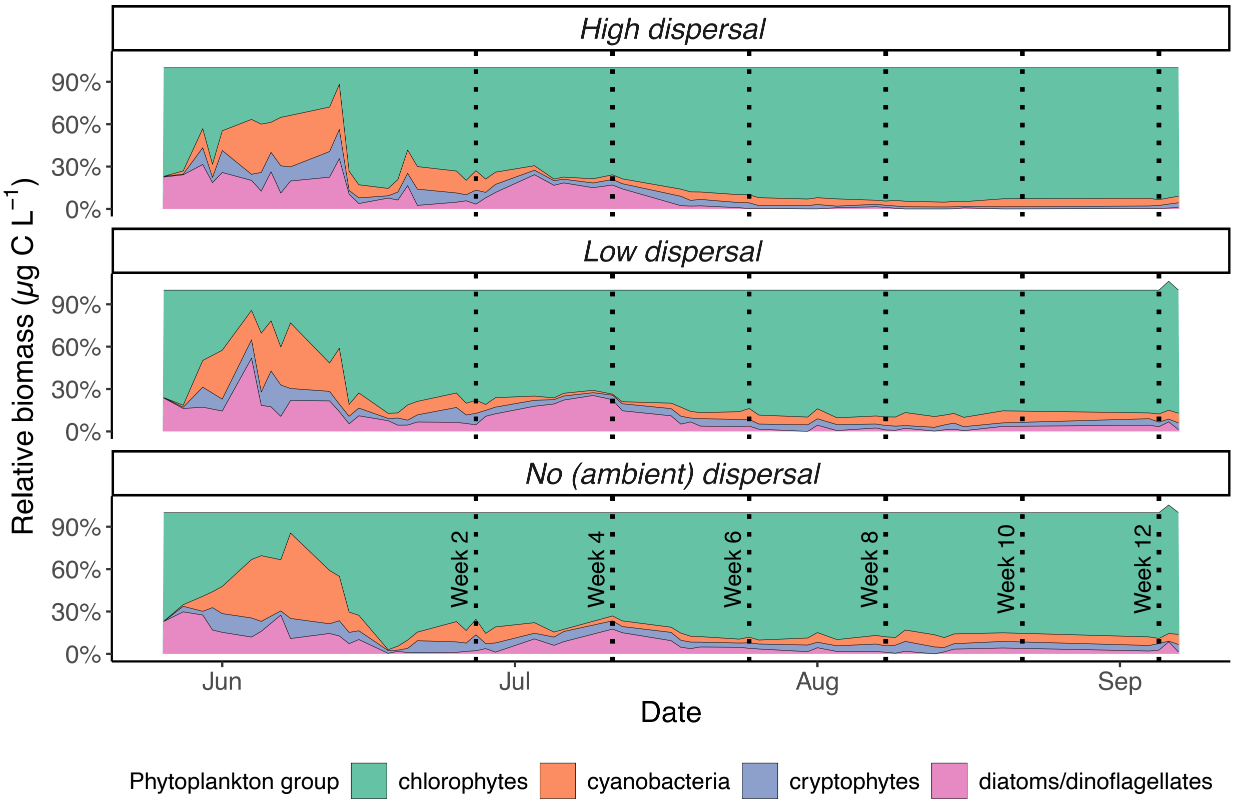


**Figure S7:** Dynamics in the relative biomass contribution of four major phytoplankton taxonomic groups by dispersal treatment. Vertical dotted lines indicate the six sampling days. Mesocosm heaters were powered on at the beginning of the time series. The initial dispersal treatment was applied two weeks before the first sampling point (week 2) and subsequently every week.

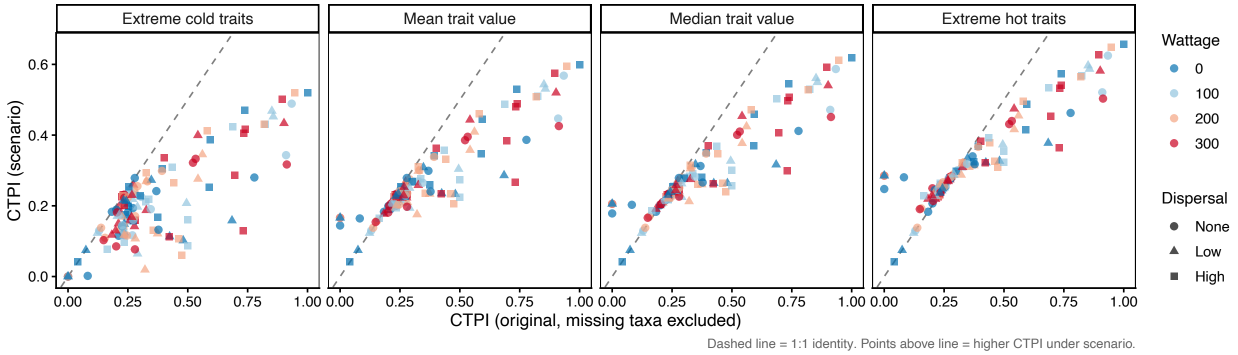


**Figure S8:** Sensitivity of Community Thermal Phenotype Index (CTPI) to different thermal trait assignments for taxa with unavailable trait data. The x-axis shows CTPI computed using only the 11 phytoplankton taxa with available published thermal performance curve parameters used in our analyses. The y-axis shows CTPI recomputed after assigning the four missing taxa an arbitrary thermal optimum ($T_{\text{opt}}$) value spanning the range observed among present species: extreme cold (16 °C), mean (25.0 °C), median (26.9 °C), and extreme hot (31 °C). Each point represents one mesocosm at one sampling time point.

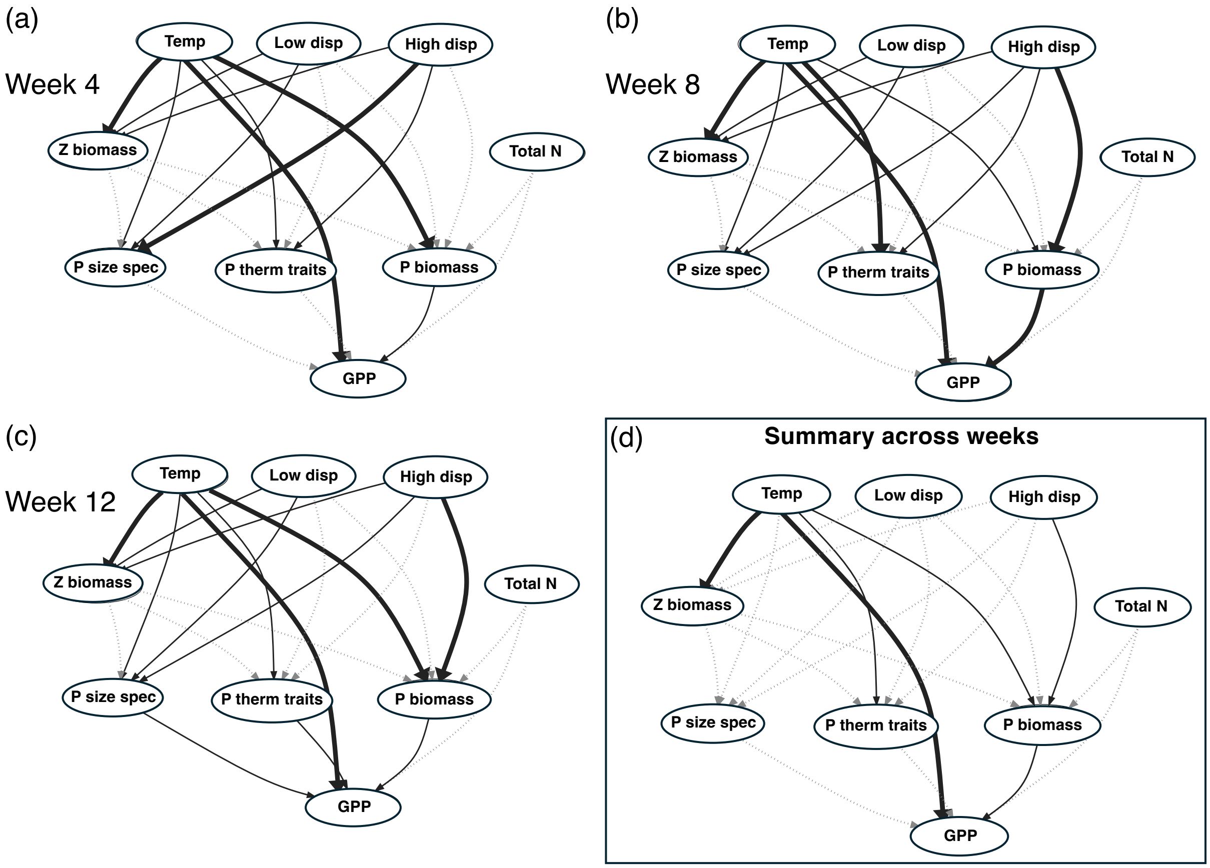


**Figure S9:** Directed acyclic graphs summarizing direct and indirect effects on GPP across all weeks based on the Structural Equation Model described in Table S2. Arrows represent all paths tested. Note Z means zooplankton, P means phytoplankton, and N means nitrogen. **(a-c)** Summary of non-zero path effects in weeks 4, 8, and 12. Thick solid arrows indicate paths that had non-zero effects at a strict ROPE width (20% of SD(GPP)). Thin solid arrows indicate paths that were not significant at ROPE = 20%, but significant at ROPE = 5% or 10% (i.e., sensitive to ROPE width). Grey dotted arrows indicate paths coefficients that are indistinguishable from zero at all ROPE widths. **(d)** Summary of paths that have non-zero effects in all three weeks. Thick solid arrows indicate paths that were non-zero at a strict ROPE width (20% of SD(GPP)) at all three time points. Thin solid arrows indicate paths that were not consistently significant at ROPE = 20%, but were significant at ROPE = 5% or 10% at all three time points. Grey dotted arrows indicate paths that did not have a consistent effect at all three time points. All posterior path coefficient estimates and % of HDI within ROPE are reported in Table S3. Posterior path coefficient sensitivities to ROPE width are reported in Table S4.

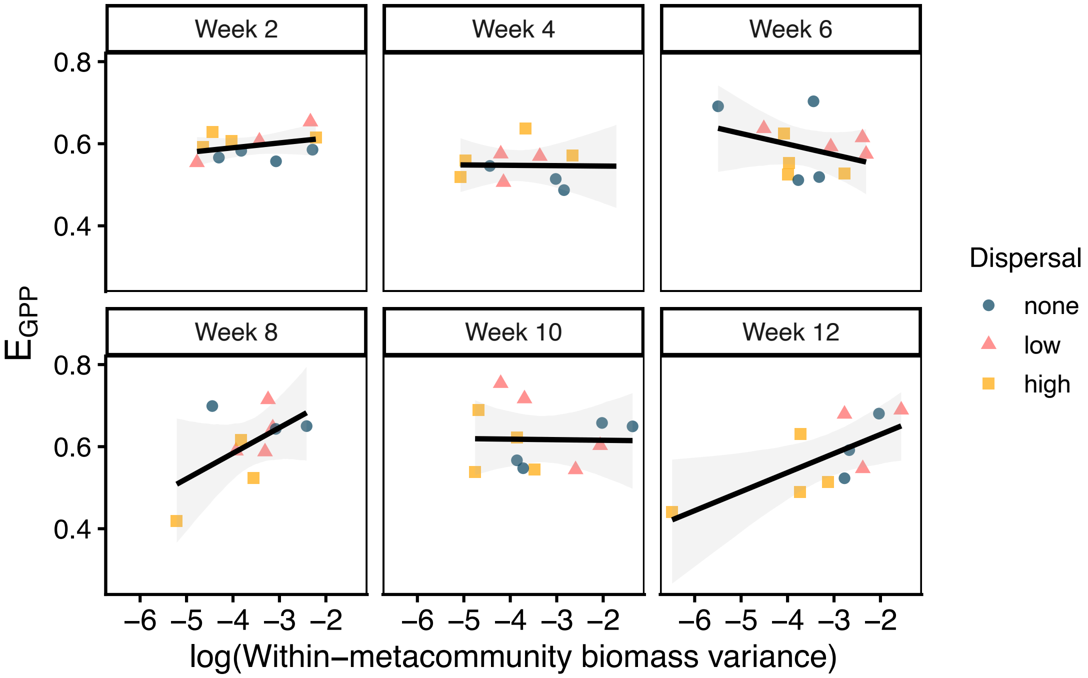


**Figure S10:** Correlation between within-metacommunity thermal sensitivity of GPP ($E_{\mathrm{GPP}}$ and within-metacommunity coefficient of variation for biomass (CV). Each point represents $E_{\mathrm{GPP}}$ estimated for one single metacommunity (n = 4 mesocosms, one of each temperature treatment) regressed against the estimated biomass CV for that community.

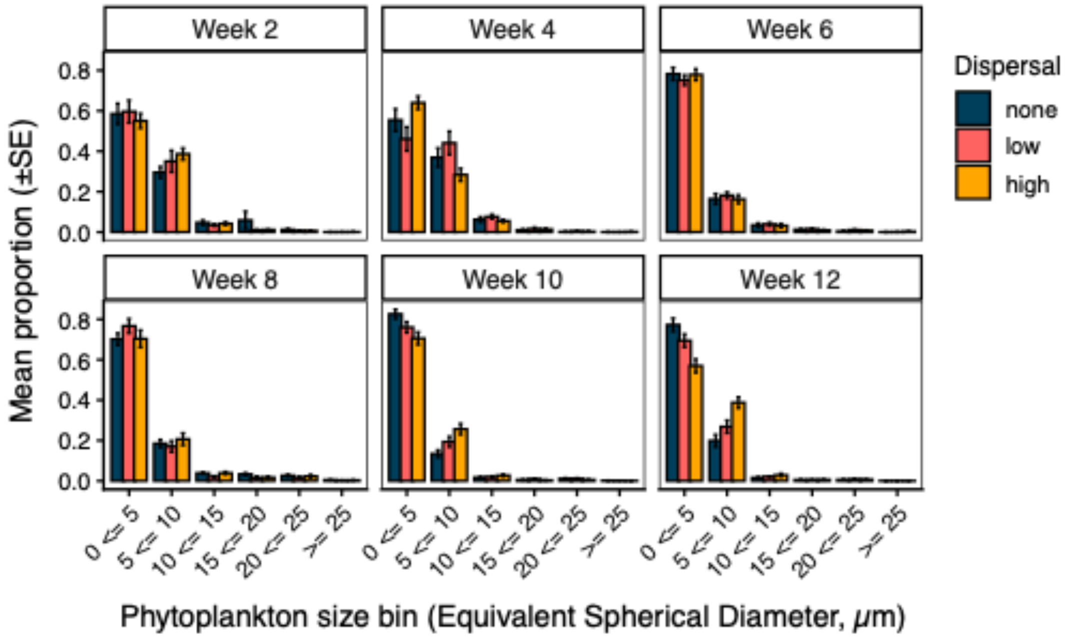


**Figure S11:** Mean proportion of phytoplankton counts (± standard error) from each 5 $\mu$m size bin by week and dispersal treatment.

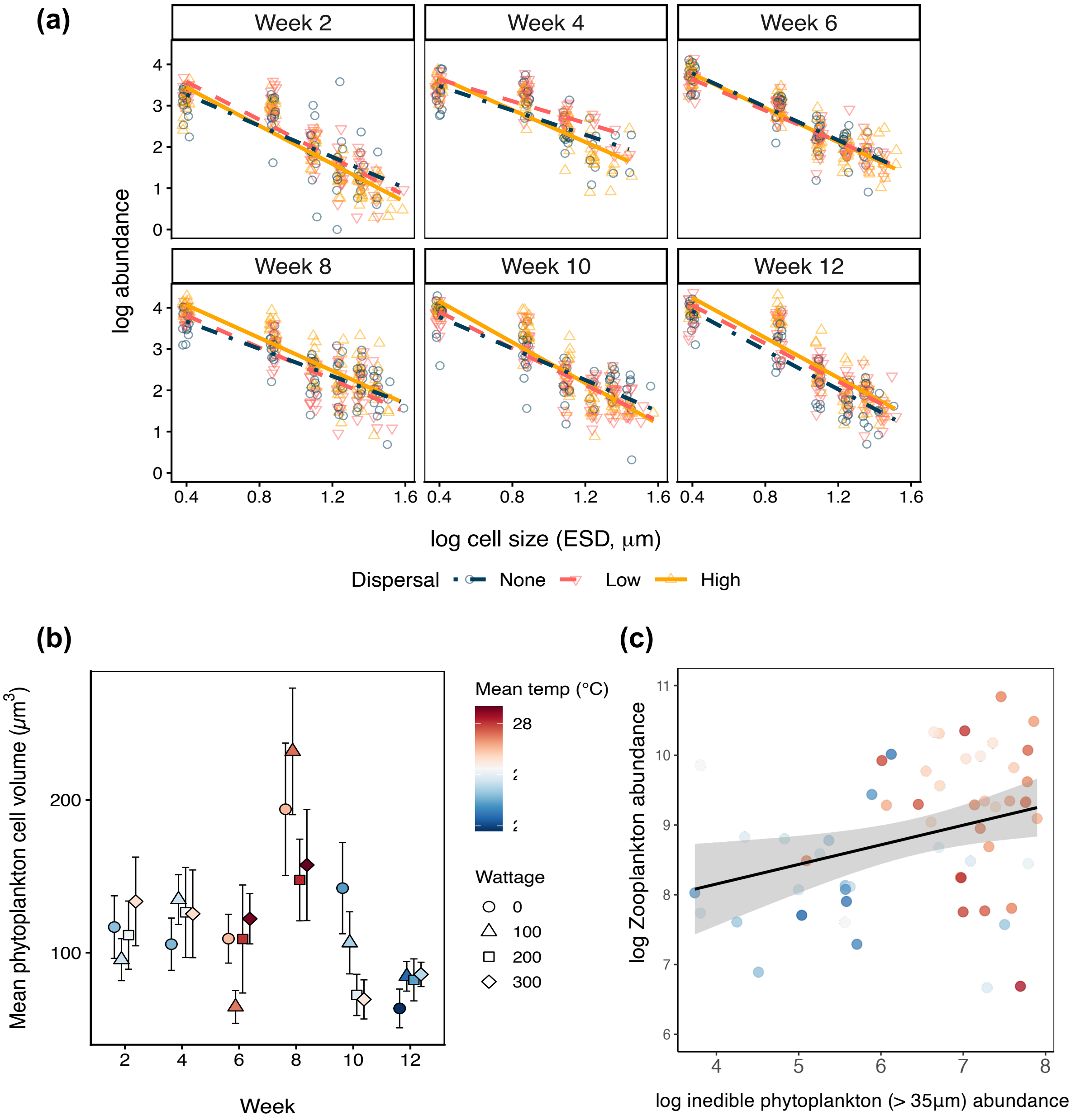


**Figure S12:** Dispersal and temperature effects on phytoplankton community size spectra and mean size. **(a)** Community size spectra by dispersal treatment across weeks. **(b)** Mean phytoplankton cell size ($\mu$g C cell$^{-1}$) over time by warming treatment. Colours indicate mean daily temperature across all mesocosms of a given warming treatment in a given week. **(c)** Temperature-mediated trophic effects on phytoplankton size over time: Zooplankton abundance versus large phytoplankton (equivalent spherical diameter >30 $\mu$m) abundance. Point colours indicate mesocosm temperature at the time of sampling.

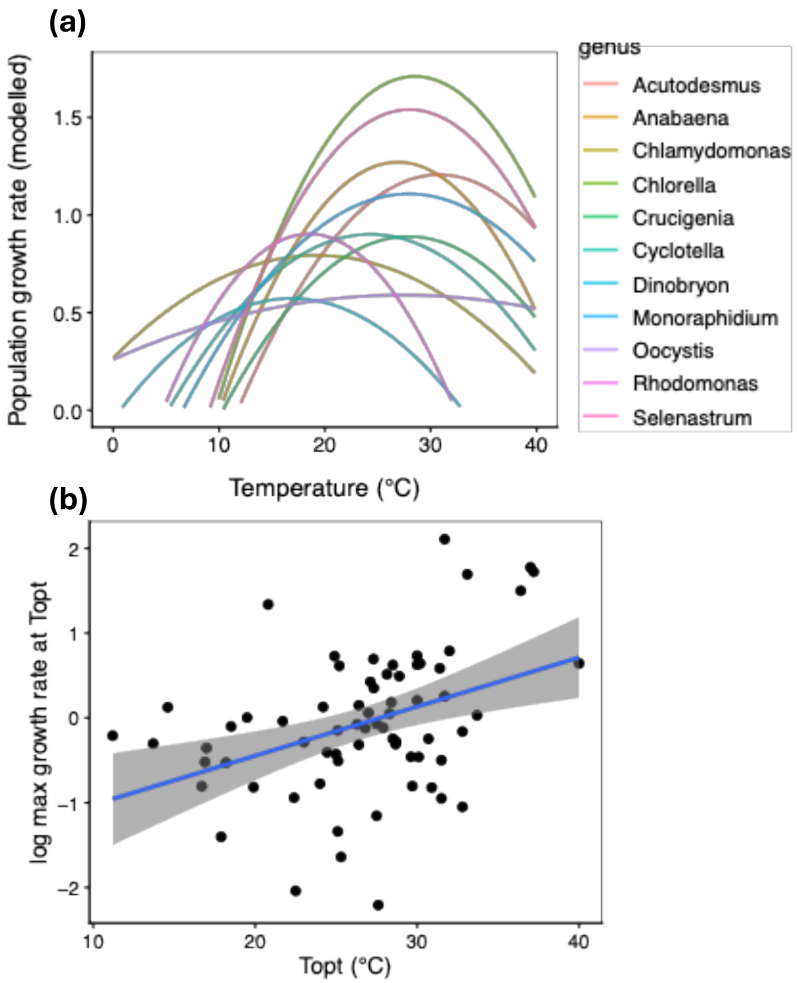


**Figure S13:** Thermal phenotypes of phytoplankton taxa found in the mesocosm experiment. Thermal response curve parameters, including $T_{\text{opt}}$ and maximum growth rate are from the published database in Thomas *et al.* 2016. **(a)** Predicted thermal response curves for all genera in the experiment based on published parameters for the Norberg model of temperature-dependent growth (Norberg, 2004). **(b)** Log-linear relationship between species' thermal optima and maximum population growth rate (day$^{-1}$. A positive relationship between $T_{\text{opt}}$ and maximum growth is a pre-requisite for a "hotter-is-better" temperature dependence of GPP across a spatial thermal gradient.

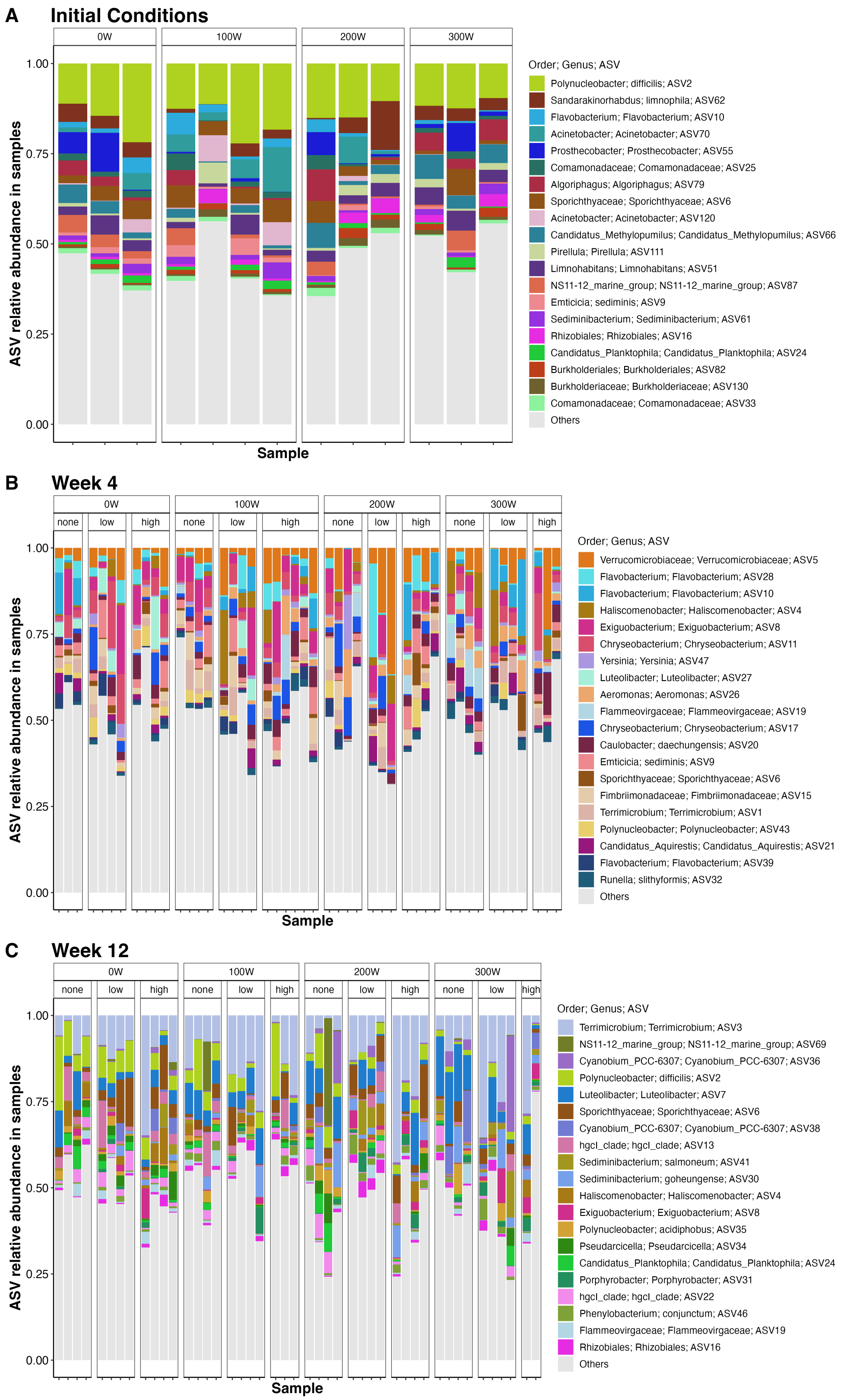


**Figure S14:** Bacterial taxonomic composition (16S) across experimental treatments in **(a)** Week 0 (initial conditions immediately before first dispersal treatment), **(b)** Week 4, and **(c)** Week 12. Colours represent the top 20 most abundant bacterial amplicon sequence variants (ASVs) at each time point.

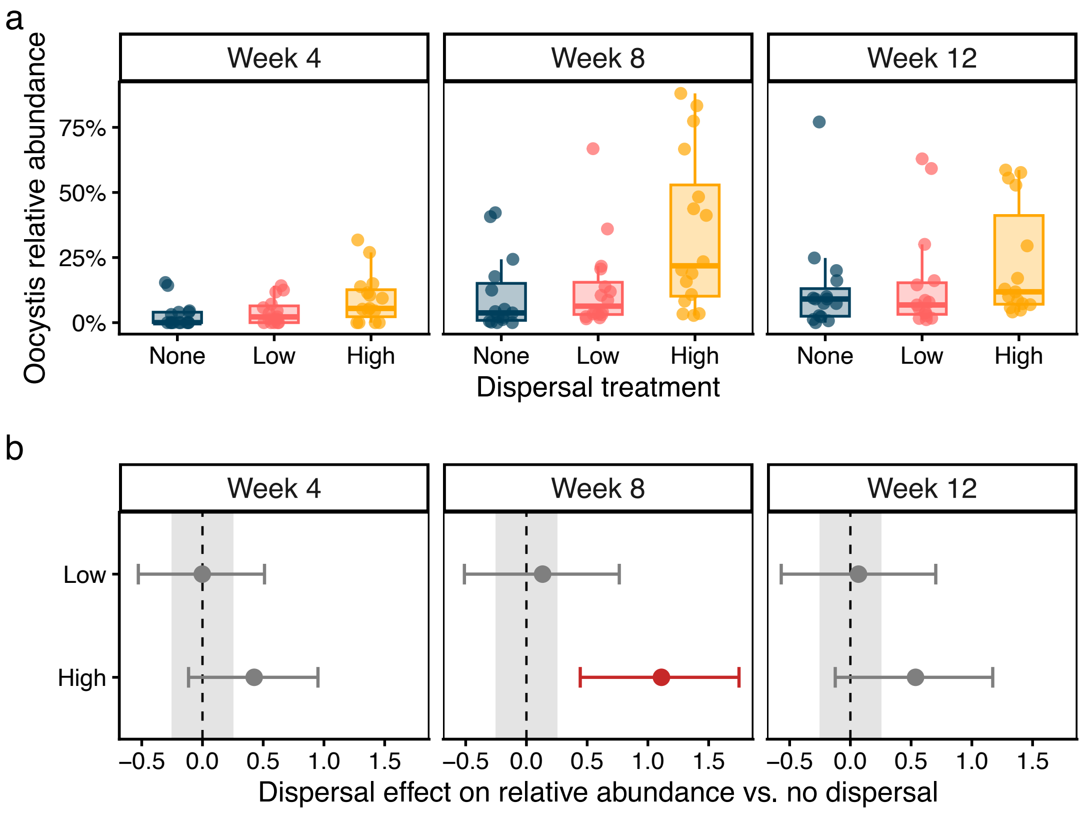


**Figure S15:** **(a)** Relative abundance of *Oocystis* sp. by dispersal treatment and week. **(b)** Posterior estimates of dispersal effect on *Oocystis* relative abundance by week, described by parameter $\beta_{D_l \times W_k}$ in the LMM described in eqn S15.

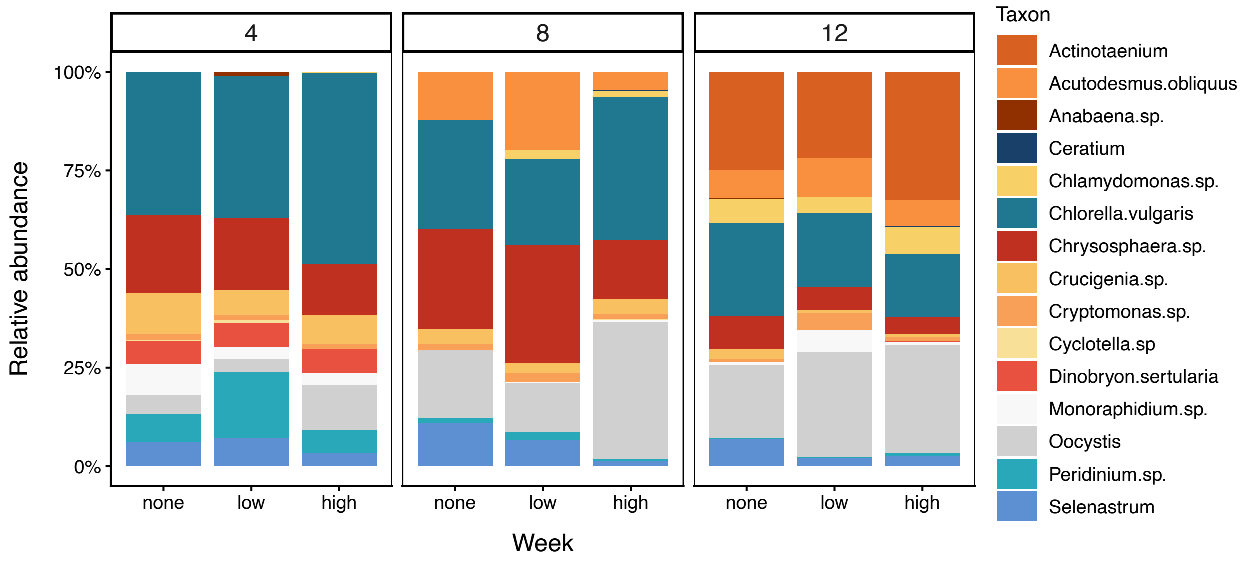


**Figure S16:** Mean relative abundances of phytoplankton taxa by dispersal treatment in weeks 4, 8, and 12.
